# Intrinsic reward-like dopamine and acetylcholine dynamics in striatum

**DOI:** 10.1101/2022.09.09.507300

**Authors:** Anne C. Krok, Pratik Mistry, Yulong Li, Nicolas X. Tritsch

**Affiliations:** Neuroscience Institute, New York University Grossman School of Medicine; New York, NY 10016, U.S.A., and Fresco Institute for Parkinson’s and Movement Disorders, New York University Langone Health, New York, NY 10016, U.S.A; State Key Laboratory of Membrane Biology, Peking University School of Life Sciences, Beijing, China

## Abstract

External rewards like food and money are potent modifiers of behavior^1,2^. Pioneering studies established that these salient sensory stimuli briefly interrupt the tonic cell-autonomous discharge of neurons that produce the neuromodulators dopamine (DA) and acetylcholine (ACh): midbrain DA neurons (DANs) fire a burst of action potentials that broadly elevates DA levels in striatum^3-5^ at the same time as striatal cholinergic interneurons (CINs) produce a characteristic pause in firing^6-8^. These phasic responses are thought to create unique, temporally-limited conditions that motivate action and promote learning^9-14^. However, the dynamics of DA and ACh outside explicitly-rewarded situations remain poorly understood. Here we show that extracellular levels of DA and ACh fluctuate spontaneously in the striatum of mice and maintain the same temporal relationship as that evoked by reward. We show that this neuromodulatory coordination does not arise from direct interactions between DA and ACh within striatum. Periodic fluctuations in ACh are instead controlled by glutamatergic afferents, which act to locally synchronize spiking of striatal cholinergic interneurons. Together, our findings reveal that striatal neuromodulatory dynamics are autonomously organized by distributed extra-striatal afferents across behavioral contexts. The dominance of intrinsic reward-like rhythms in DA and ACh offers novel insights for explaining how reward-associated neural dynamics emerge and how the brain motivates action and promotes learning from within.

## Main Text

Rewards and the sensory cues that predict their availability elicit brief changes in the release of the neuromodulators dopamine (DA) and acetylcholine (ACh). These phasic responses are thought to stand out from the basal, steady concentration of DA and ACh that brain circuits normally bathe in to affect behavior. In the mammalian striatum, rewards evoke a phasic increase in extracellular DA^3-5^ and a phasic decrease in ACh^6-8^, the coincidence of which is critical for DA to act as a teaching signal, as muscarinic M4 receptors oppose DA-dependent synaptic plasticity on direct-pathway striatal projection neurons (SPNs)^15-17^. Recent studies have also implicated phasic elevations in DA in the initiation and invigoration of self-paced movements^18-21^, and ACh has been proposed to help striatal circuits distinguish between DA signals related to motor performance and learning^12-14,22,23^. However, the dynamics of extracellular DA and ACh levels in vivo remain unknown. Because DA and ACh interact extensively in striatum without necessarily engaging somatic spiking^24-30^, addressing this question requires approaches that directly and simultaneously report DA and ACh levels on sub-second time scales.

To reveal the dynamics of striatal DA and ACh, we imaged the D2 receptor-based red fluorescent DA indicator rDA1m^31^ and the muscarinic M3 receptor-based green fluorescent ACh indicator ACh3.0^32^ concurrently from the dorsolateral striatum (DLS) of mice head-fixed on a cylindrical treadmill using fiber photometry (**Fig. 1a,b**; **Extended Data Fig. 1a,b**). Mice spontaneously alternated between bouts of immobility and locomotion, and on a subset of recording sessions were provided with uncued liquid rewards. As expected, rewards evoked phasic changes in the fluorescence of both sensors reminiscent of the characteristic discharge of midbrain DANs^3-5^ and striatal CINs^6-8^ (**Fig. 1c-e**). On average, rDA1m showed a sharp increase in fluorescence (mean amplitude: 9.9 ± 1.0 % ΔF/F; n = 13 mice) that decayed back to baseline within 1.5 s, whereas ACh3.0 showed a phasic dip in fluorescence below a stable baseline (mean amplitude: 4.4 ± 0.5 % ΔF/F; mean duration: 298 ± 21 ms) preceded by a brief increase in fluorescence (mean amplitude: 2.9 ± 0.7 % ΔF/F). These events appeared in a sequence, starting with the ACh3.0 peak followed 82 ± 5 ms later by the rDA1m peak, itself followed by the dip in ACh3.0 fluorescence (time to trough: 200 ± 17 ms) (**Fig. 1f,g**). These data are consistent with the sequence of reward-evoked action potential responses observed from putative DANs and CINs in rodents and primates, confirming our ability to detect and isolate phasic changes in striatal DA and ACh from simultaneous photometric recordings.

**Fig. 1.**
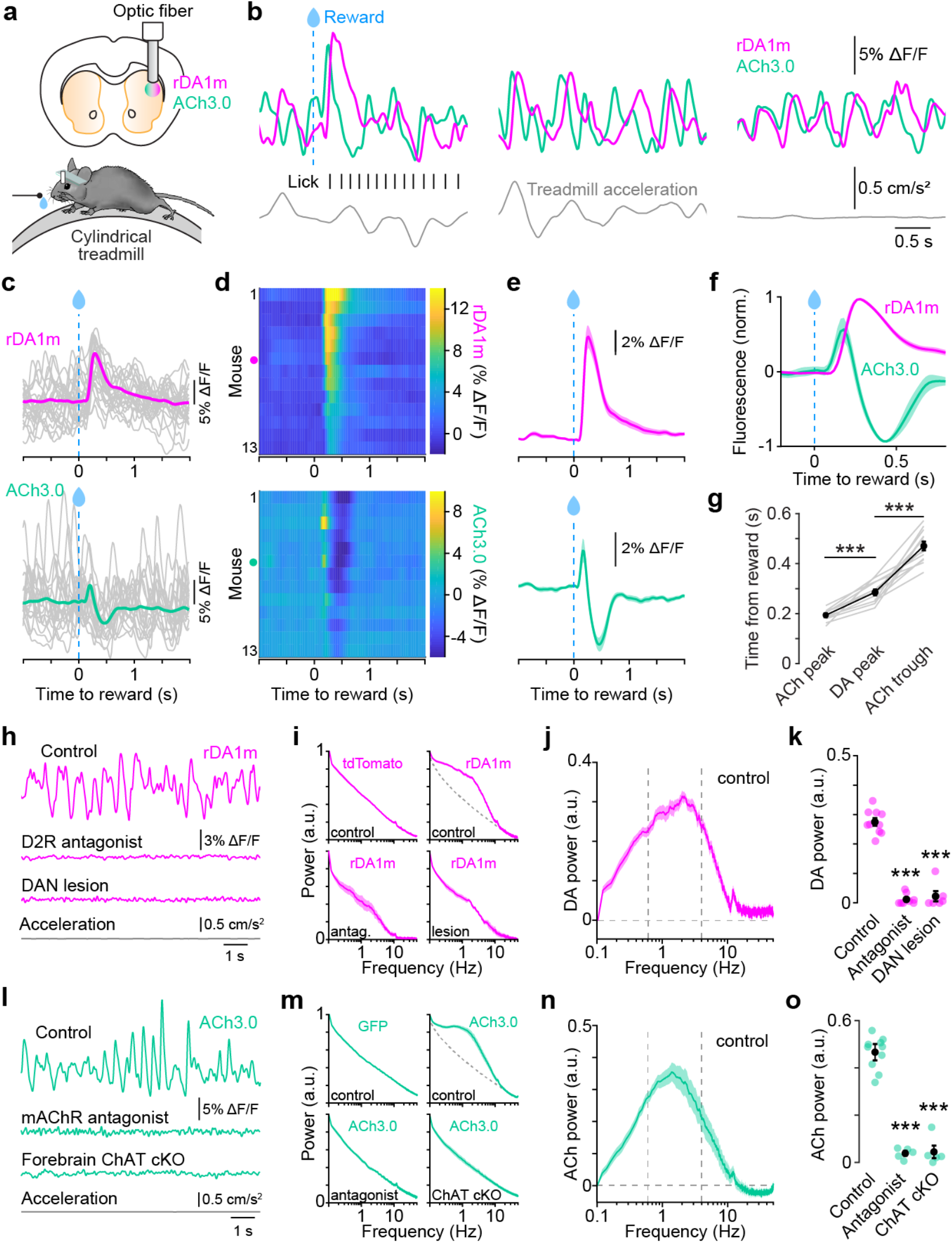
Spontaneous periodic fluctuations in striatal DA and ACh levels. **a**, Schematic of fiber photometry imaging of rDA1m and ACh3.0 fluorescent sensors in DLS (*top*) and experimental setup (*bottom*). **b**, Example rDA1m (magenta) and ACh3.0 (teal) fluorescence signals (*top*) recorded simultaneously during reward (*left*), locomotion (*middle*) and immobility (*right*), with water delivery (blue line), lick events (black) and treadmill acceleration (gray). **c**, *Top*: single imaging session-averaged rDA1m fluorescence aligned to reward delivery (magenta) with 20 randomly-selected single trials overlaid in gray. *Bottom*: same for ACh3.0 in the same session. **d**, Heatmap of rD1mA (*top*) and ACh3.0 (*bottom*) fluorescence aligned to reward delivery for all mice. Dots indicate mouse in (**c**). **e**, Group-averaged DA (*top*) and ACh (*bottom*) fluorescence aligned to reward delivery (n = 13 mice). **f**, Same as **e** with DA and ACh signals normalized to respective maxima and minima. **g**, Latency of reward-evoked DA and ACh transients. Individual mice shown in gray, population mean (± sem) in black. ACh peak vs. DA peak: p = 2.0×10^−7^; DA peak vs. ACh trough: p = 9.6×10^−10^; Dunn’s multiple comparisons. **h**, Example DA fluorescence traces during immobility under control conditions (*top*), following systemic administration of the D2R antagonist sulpiride (*middle*) or 6OHDA-mediated lesioning of midbrain DA neurons (*bottom*). Acceleration (gray) shown for control recording. **i**, Baseline-adjusted and normalized power spectra of raw photometry signal for tdTomato (n = 3) and rDA1m under control conditions (n = 13), in D2R blocker (n = 8) or following 6OHDA lesion (n = 6) during periods of immobility. **j**, Power spectrum of DA signal isolated by subtracting the normalized power spectrum of tdTomato from the control in **i** during periods of immobility.Vertical lines indicate 0.5 and 4 Hz. **k**, DA power in 0.5-4 Hz frequency band for control mice, in D2R blocker (p = 1.0×10^−10^) or following 6OHDA lesion (p = 2.4×10^−8^), both vs. control, Student’s two-sample *t*-test. Population mean (± sem) shown in black. **l**, Same as **h** for ACh3.0 under control conditions (*top*), following systemic administration of a mAChR antagonist (*middle*), and in ChAT cKO mice (*bottom*). **m**, Same as **i** for GFP (n = 3) and ACh3.0 (n = 13 controls, 5 antagonists, 5 ChAT cKO). **n**, Same as **j** for ACh power. **o**, Same as **k** for ACh power. Antagonist: p = 1.7×10^−8^; cKO: p = 4.6×10^−8^; both vs. control, Student’s two-sample *t*-test.

### Striatal DA and ACh levels fluctuate spontaneously

rDA1m and ACh3.0 fluorescence signals also showed large, periodic fluctuations outside of rewards, including during periods of immobility and in the absence of overt sensory stimuli that were not apparent in session-averaged and reward-aligned data (**Fig. 1b, h, l; Extended Data Fig. 1c-g**). These spontaneous signal fluctuations, which occurred most prominently in the 0.5–4 Hz frequency band (mode for DA: 2.0 ± 0.2 Hz; ACh: 1.8 ± 0.2 Hz; **Fig. 1j, n**) were also observed before mice ever received rewards on the treadmill, during periods of immobility and active locomotion (**Extended Data Fig. 1f, g**), as well as in an open field arena (**Extended Data Fig. 2**), but not in mice expressing green fluorescent protein (GFP) or tdTomato (**Fig. 1i, m**), confirming that they do not reflect hemodynamic, movement, or signal processing artifacts. In mice expressing rDA1m alone, spontaneous fluorescence transients during immobility were reversibly blocked following systemic administration of the D2 receptor (D2R) antagonist sulpiride (n = 8 mice) and were entirely absent following infusion of the toxin 6-hydroxydopamine (6-OHDA) in the substantia nigra pars compacta (n = 6 mice; **Fig. 1h, i, k**), showing that rDA1m fluorescence transients in DLS reflect DA release from midbrain DANs. In ACh3.0-expressing mice, spontaneous fluorescent transients disappeared upon treatment with the muscarinic ACh receptor (mAChR) antagonist scopolamine (n = 5 mice) and were not detected in mice in which the ACh synthetic enzyme choline acetyltransferase (ChAT) is conditionally knocked out from forebrain cholinergic neurons (ChAT^f/f^;Nkx2.1^Cre^ mice, referred to as ChAT cKO; n = 5 mice; **Fig. 1l-o**), indicating that ACh3.0 fluctuations reflect spontaneous changes in ACh release from local CINs and not from brainstem cholinergic afferents^33^. Together, these data show that striatal circuits constantly undergo large, periodic increases and decreases in extracellular DA and ACh levels originating from midbrain DANs and striatal CINs, respectively.

### DA and ACh transients maintain a consistent temporal relationship across behavioral states

Striatal neurons co-express receptors for DA and ACh that engage competing intracellular signaling pathways^34,35^. The net impact of phasic changes in DA and ACh levels on striatal function therefore depends on their relative timing and amplitude. During rewards, DA and ACh were negatively correlated (Pearson correlation coefficient r = –0.48 ± 0.02, n = 13 mice) with a temporal lag of 123 ± 9 ms (**Fig. 2a-c**), indicating that peaks in DA slightly precede dips in ACh. Importantly, DA and ACh showed a similar negative correlation outside of reward, whether mice were actively locomoting or remained immobile (**Fig. 2a-c**). In agreement with this, we observed strong coherence between DA and ACh in the 0.5–4 Hz frequency band (**Fig. 2d, e**) that maintained a phase offset of approximately 90 degrees throughout our recordings and across all conditions (**Fig. 2f-l**). The amplitude of individual DA peaks occurring coincidentally with dips in ACh also showed considerable overlap during reward, locomotion and immobility (**Extended Data Fig. 1h, i**), further highlighting difficulties in distinguishing these events categorically. These data therefore show that DA and ACh transients maintain a consistent temporal relationship across behavioral states and suggest that reward and other salient stimuli do not create unique neuromodulatory dynamics in striatum, but instead recruit intrinsically-structured rhythms in DA and ACh.

**Fig. 2.**
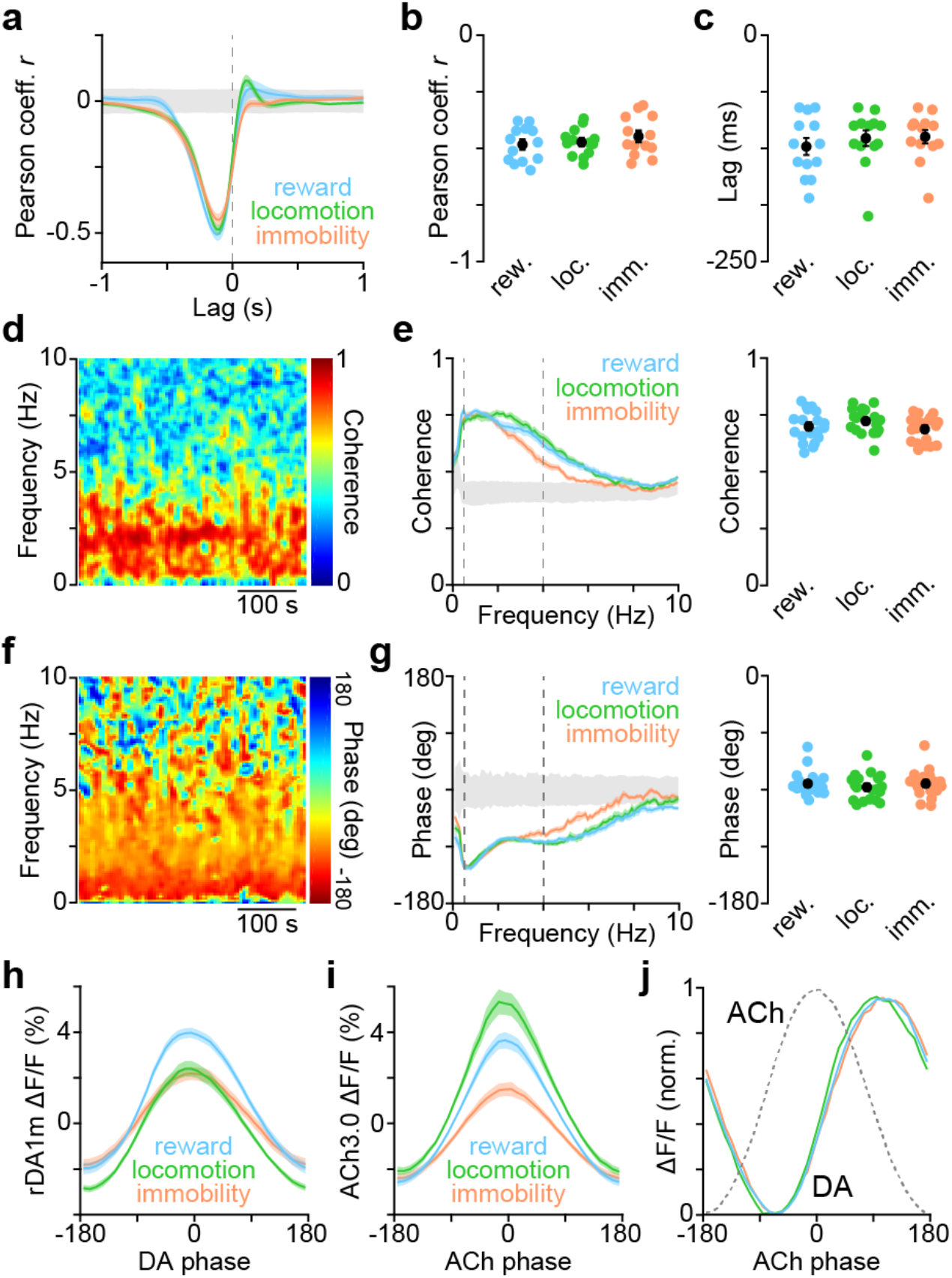
DA and ACh maintain a consistent temporal relationship across behavioral states. **a**, Mean (± sem) cross-correlation between simultaneously recorded DA and ACh signals (n = 13 mice) during reward (blue), locomotion (green) and immobility (orange) with 95% confidence interval (gray). **b**, Peak correlation coefficient (Pearson’s *r*) between DA and ACh signals in individual mice (n = 13) across behavioral states. Population mean (± sem) shown in black. p = 0.5, one-way ANOVA. **c**, Same as (**b**) for time lag of negative cross-correlation peak. p = 0.6, one-way ANOVA. **d**, Magnitude of coherence between DA and ACh signals across frequency and time domains for example recording. **e**, *Left*, mean (± sem) coherence between DA and ACh signals at different frequencies across behavioral states (n = 13). Vertical lines depict 0.5 and 4 Hz. *Right*, median coherence at 0.5-4 Hz for individual mice. Population mean (± sem) in black. p = 0.06, one-way ANOVA. **f**, Same as **d** for phase offset between DA and ACh. **g**, Same as **e** for phase offset between DA and ACh. p = 0.4, one-way ANOVA. **h**, Mean (± sem) DA fluorescence (n = 13) at different phases of periodic DA fluctuations in the 0.5-4 Hz frequency band during reward (blue), locomotion (green) and immobility (orange). **i**, Same as **h** for ACh fluorescence vs. phase of periodic ACh fluctuations. **j**, Peak-normalized DA fluorescence vs. phase of periodic ACh fluctuations (shown as gray dotted line) in 0.5-4 Hz frequency band during reward (blue), locomotion (green) and immobility (orange).

### DA and ACh fluctuations are locally coherent

To determine whether stereotyped coupling between DA and ACh extends to other striatal regions, we co-expressed and simultaneously imaged rDA1m and ACh3.0 in DLS as well as in dorsomedial striatum (DMS). Within DMS, extracellular DA and ACh levels fluctuated constantly and coherently at 0.5–4 Hz (**Extended Data Fig. 3a-f**), and were consistently phase-shifted irrespective of behavioral state (**Extended Data Fig. 3g-j**), showing that DA and ACh maintain a similar temporal relationship in DMS. To determine if these fluctuations occur concurrently throughout dorsal striatum, we compared DA transients in one striatal region to ACh signals in the other. DA and ACh signals in DLS and DMS were not coherent (**Extended Data Fig. 3k, l**), indicating that DA and ACh are not entrained by global brain-wide rhythms like breathing. Thus, although DA and ACh are stereotypically coupled locally, their coherence declines with distance, such that periodic fluctuations in DLS occur largely independently from those in DMS.

### Periodic DA and ACh fluctuations are not coordinated locally within striatum

Local coordination between DA and ACh may result from direct interactions between these neuromodulators within striatum. Experiments in brain slices convincingly demonstrated that DA release can briefly pause CIN firing via D2 receptors ^29,30^, while synchronous activation of CINs can evoke action potentials in DAN axons and cause widespread release of DA in striatum following the activation of presynaptic β2-containing nicotinic ACh receptors (nAChRs)^24-28^. To investigate whether DA locally entrains ACh release in vivo, we simultaneously imaged DA and ACh in DLS before and after locally infusing a control saline solution or a cocktail of DA receptor antagonists (n = 4 mice; **Fig. 3a**). Within a few minutes of infusion, the latter effectively prevented the rDA1m sensor from reporting spontaneous and reward-evoked changes in extracellular DA levels (**Fig. 3b, c**), confirming our ability to block DA receptor signaling in the area imaged by our fiber optic. By contrast, this manipulation did not alter the timing or magnitude of spontaneous or reward-evoked ACh fluctuations compared to saline-infused mice (**Fig. 3d-f; Extended Data Fig. 4a-c**). To exclude contributions from other transmitters co-released by DANs, including glutamate and GABA ^30,36^, we monitored ACh in DLS before and after lesioning midbrain DANs with 6-OHDA (n = 6 mice; **Extended Data Fig. 4d**); ACh levels continued to show periodic fluctuations during immobility that were indistinguishable from those observed in the same animals prior to lesion (**Extended Data Fig. 4e-h**).

**Fig. 3.**
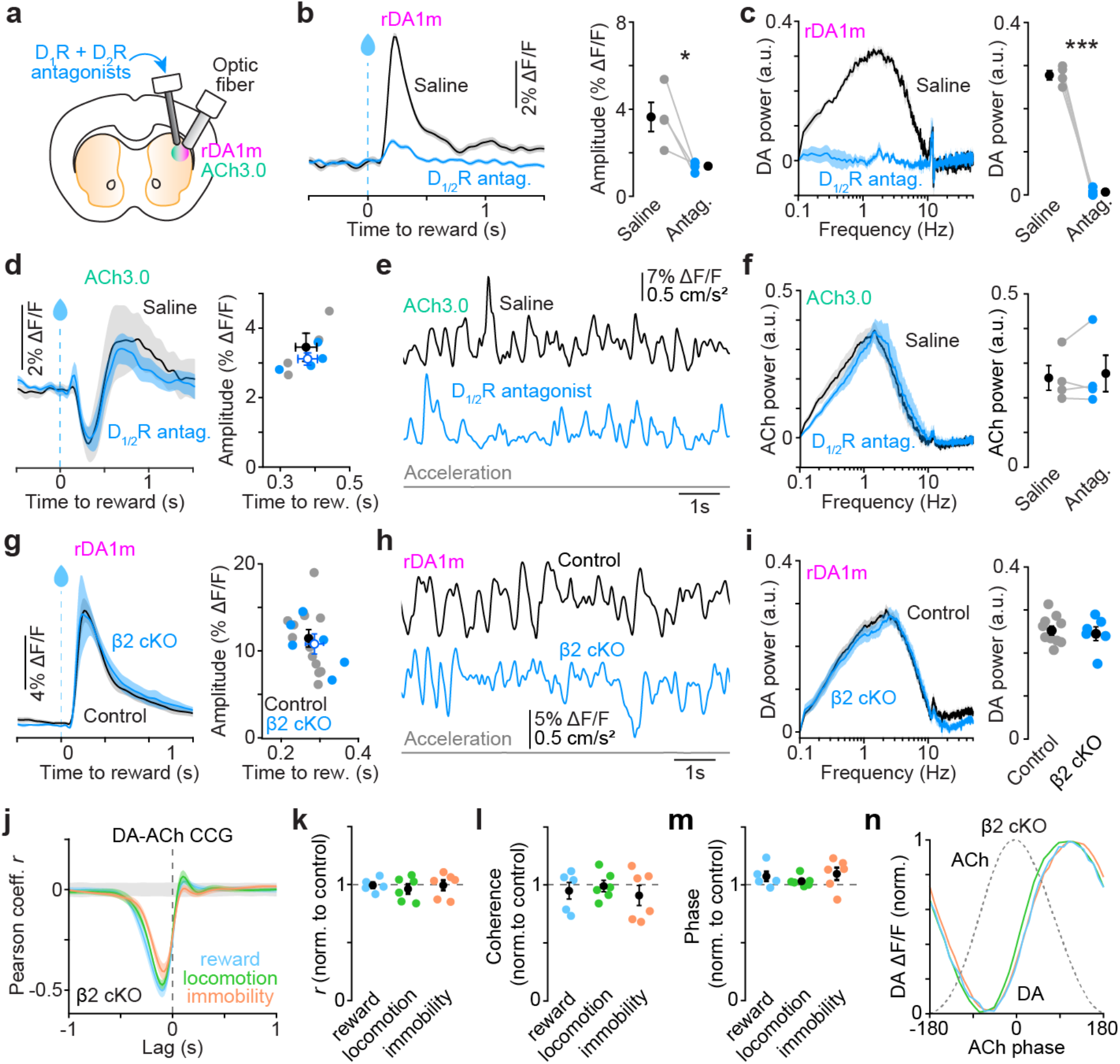
Periodic fluctuations in DA and ACh are not coordinated locally within striatum. **a**, Experimental preparation for local pharmacological inhibition of DA signaling in DLS. **b**, *Left*, example imaging session: DA fluorescence aligned to reward delivery after intra-striatal infusion of saline or a cocktail of D1R and D2R antagonists. *Right*, reward-evoked DA amplitude after DLS infusion of saline and D1/2R antagonists (n = 4 mice). Population mean (± sem) shown in black. p = 0.03 vs. saline, Student’s paired *t*-test. **c**, *Left*, power spectrum of DA signal during immobility after DLS infusion of saline or D1/2R antagonists. *Right*, mean (± sem) DA power at 0.5-4 Hz; p = 3.31×10^−4^, Student’s paired *t*-test. **d**, *Left*, same as **b** for ACh. *Right*, scatter plot of reward-evoked ACh dip amplitude and timing in n = 4 mice infused with saline (gray) and D1/2R antagonists (blue). Amplitude, p = 0.5; timing, p = 0.9, both vs. saline, Student’s paired *t*-test. **e**, Example ACh fluorescence traces after DLS infusion of saline or D1/2R antagonist during immobility. Treadmill acceleration for D1/2R antagonist shown in gray. **f**, Same as (**c**) for ACh power. p = 0.4 vs. saline, Student’s paired *t*-test. **g**, *Left*, example reward-evoked DA transients in control (black) and β2 cKO (blue) mice. *Right*, scatter plot of reward-evoked DA peak amplitude and timing in control (n = 13; gray) and β2 cKO mice (n = 6; blue). Group means (± sem) shown in black. Amplitude, p = 0.08; timing, p = 0.5, both vs. saline, Student’s two-sample *t*-test. **h**, Example DA fluorescence traces from a control (black) and β2 cKO mouse (blue) during immobility. Acceleration of β2 cKO mouse shown in gray **i**, Same as **c** for DA power in control vs. β2 cKO mice. p = 0.7, Student’s two-sample *t*-test. **j**, Mean (± sem) DA–ACh cross-correlogram (CCG) from β2 cKO mice (n = 6) during reward (blue), locomotion (green) and immobility (orange). **k**, Pearson correlation coefficient *r* in β2 cKO mice normalized to control during reward (blue; p = 0.6), locomotion (green; p = 0.9) and immobility (orange; p = 0.4). Group mean (± sem) shown in black. All comparisons vs. control mice, Student’s two-sample *t*-test. **l**, Same as **k** for median DA–ACh coherence in 0.5-4Hz band (reward: p = 0.9; locomotion: p = 0.3; immobility: p = 0.9). **m**, Same as **k** for median phase offset (reward: p = 0.4; locomotion: p = 0.6; immobility: p = 0.2). **n**, Peak-normalized DA fluorescence in β2 cKO mice vs. phase of periodic ACh fluctuations (shown as gray dotted line) in 0.5-4 Hz frequency band during reward (blue), locomotion (green) and immobility (orange).

Next, we investigated whether ACh might directly entrain DA release within striatum. We generated mice in which CINs cannot directly evoke axonal release of DA by conditionally deleting the β2 nAChR subunit specifically in DANs (Dat^IRES-Cre/+^;β2^fl/fl^ mice; referred to as β2 cKO) and imaged ACh3.0 and rDA1m simultaneously in DLS using photometry (**Extended Data Fig. 5a**). β2 cKO mice had intact ACh and DA responses to uncued rewards (**Fig. 3d; Extended Data Fig. 5b**), confirming that striatal DA transients triggered by reward derive from the well-established somatic burst response of midbrain DANs. Outside rewards, DA and ACh transients were similarly unperturbed, fluctuating periodically at the same rate (**Fig. 3h, i; Extended Data Fig. 5c, d**) and maintaining their strong coherence and phase relationship during locomotion as well as immobility (**Fig. 3j-n; Extended Data Fig. 5e, f**).

To exclude the possibility that homeostatic adaptations occlude the influence that ACh normally exerts on DA terminals, we acutely blocked nAChR signaling by locally infusing the nAChR antagonist DHβE in DLS (**Extended Data Fig. 5g**). Compared to mice treated with saline, DHβE did not significantly affect spontaneous DA and ACh fluctuations (**Extended Data Fig. 5h-j**), their coherence or phase relationship during either reward, locomotion or immobility (**Extended Data Fig. 5k-m**), confirming that the strong coupling between DA and ACh is not dependent on striatal nAChR signaling in vivo. Together, these data show that the consistent temporal relationship between DA and ACh levels is not generated locally via direct molecular interactions between both neuromodulators.

### Local CIN spike coherence underlies periodic fluctuations in ACh

Our failure to reveal strong intra-striatal interactions between DA and ACh suggests that their coupling may instead be inherited from shared extra-striatal network dynamics. Previous studies indicate that phasic DA signals in striatum arise from coherent firing amongst populations of midbrain DANs ^37-41^, which integrate inputs distributed throughout the brain^42^. Striatal CINs also fire coherently in response to reward and movement^6-8,23^, but little is known about the factors that shape striatal ACh levels during immobility. To reveal how periodic ACh transients emerge, we recorded spiking activity of putative CINs (pCINs) from DLS using high-density multi-shank silicon probes. We distinguished pCINs, which make up approximately 2% of striatal neurons by established electrophysiological criteria, including spike waveform, firing rate and discharge properties (**Fig. 4a-c; Extended Data Fig. 6a, b**). Consistent with previous reports^6-8^, pCINs were tonically active (firing rate: 5.7 ± 0.1 spikes per s; n = 150 pCINs from 51 recordings in 15 mice) and exhibited irregular, non-bursting firing. Their spikes tended to occur synchronously, with 91.7% of pairs of simultaneously recorded pCINs (n = 157 pairs) showing a significant peak in their spike cross-correlogram (CCG) at a lag of 0 s (**Fig. 4d-g**). In addition, 30.6 % of pairs showed significant troughs at +0.25 and –0.25 s (**Fig. 4h**), consistent with a 2 Hz rhythm.

**Fig. 4.**
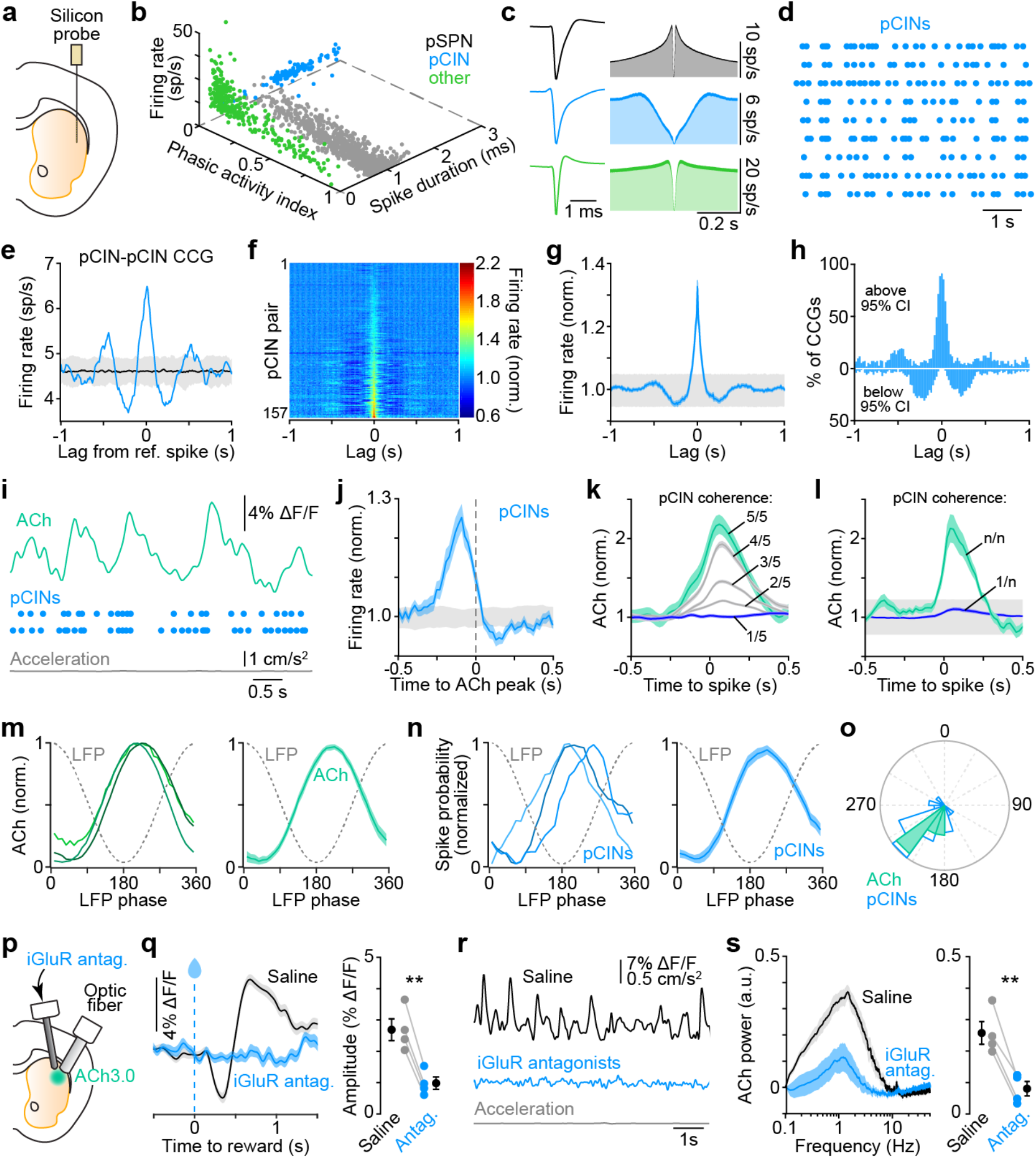
Glutamatergic inputs synchronize CIN spiking and drive periodic ACh fluctuations. **a**, Diagram of experimental configuration. **b**, Scatter plot of spike properties used to distinguish units as pSPNs (gray), pCINs (blue), or other interneurons (green). **c**, Average waveform (*left*) and auto-correlograms (*right*) for pMSNs (black), pCINs (blue), and other interneurons (green). **d**, Example spike raster for 9 simultaneously recorded pCINs during immobility. **e**, Example spike CCG for two units in **d** (blue) and 95% confidence interval (gray). **f**, Heatmap of spike CCGs for all pCIN pairs (n = 157) normalized to baseline firing rate during immobility. **g**, Mean (± sem) CCG from **f**. 95% confidence interval shown in gray. **h**, Proportion of CCGs from **f** exceeding 95% confidence interval (bin: 20 ms). **i**, Example simultaneous recording of ACh fluorescence (teal) and pCIN spiking (blue) within DLS during immobility. Treadmill acceleration in gray. **j**, Mean (± sem) firing rate of pCINs (n = 22; normalized to baseline) aligned to ACh fluorescence peaks. 95% confidence interval shown in gray. **k**, Example pCIN spike-aligned ACh fluorescence stratified based on whether 1, 2, 3, 4 or 5 out of 5 concurrently-recorded pCINs are co-active. **l**, Mean (± sem) pCIN spike-aligned ACh fluorescence when only 1 (blue) vs. all (green) concurrently-recorded pCINs are co-active, with 95% confidence interval in gray (n = 22 units). **m**, *Left*, 3 example ACh photometry recordings aligned to phase of LFP oscillation at 0.5-4 Hz (gray dotted line). *Right*, mean (± sem) peak-normalized ACh fluorescence (teal) relative to 0.5-4 Hz LFP phase. **n**, Same as **m** for pCIN spike recordings (*left*, 3 example pCINs; *right*, mean (± sem) for 94.5% (52 of 55) pCINs that showed a significant phase relationship to 0.5-4 Hz LFP. **o**, Polar plot of preferred phase of individual pCINs (blue) and ACh recordings (teal) relative to 0.5-4 Hz LFP. **p**, Diagram of experimental configuration. **q**, *Left*, example reward-evoked ACh response in a mouse infused in DLS with saline (black) or iGluR antagonists (blue). *Right*: amplitude of reward-evoked ACh trough (n = 4 mice). Group mean (± sem) shown in black. p = 0.001, Student’s paired *t*-test. **r**, Example ACh traces during immobility after saline (black) or iGluR antagonists (blue) infusion in DLS. Acceleration (gray) shown for saline. **s**, *Left*, ACh power spectrum during immobility after DLS infusion of saline or iGluR antagonists. *Right*, mean (± sem) ACh power at 0.5-4 Hz; p = 0.006, Student’s paired *t*-test.

To assess whether coordinated pCIN firing accounts for periodic fluctuations in extracellular ACh, we recorded the discharge of pCINs while simultaneously imaging ACh within overlapping regions of DLS (**Fig. 4i**). We observed a clear relationship, where phasic increases in the firing of pCINs preceded peaks in ACh, while dips in ACh fluorescence followed coordinated decreases in pCIN firing (**Fig. 4j; Extended Data Fig. 6c-i**). In both cases, fluorescence lagged pCIN firing by ∼100 ms, providing an estimate of the delay imposed by ACh3.0 sensor kinetics. To determine whether the magnitude of ACh fluctuations correlates with the degree of spike coherence amongst pCINs, we compared spike-aligned ACh signals across different periods of network synchrony using the fraction of synchronously active pCINs as a proxy. Spikes occurring during periods of high coherence (i.e. when all recorded pCINs are synchronously active) were associated with large ACh transients that gradually declined in amplitude as coherence decreased (**Fig. 4k,l**). By contrast, spikes occurring in isolation (i.e. during periods of low network coherence) did not result in appreciable changes in extracellular ACh levels. Consistent with this, ACh fluorescence increased during locomotion together with pCIN coherence (**Extended Data Fig. 7a-h**), with pCIN discharge and ACh signal peaking at maximum positive acceleration (**Extended Data Fig. 7i-k**)^23^. Importantly, ACh signals reflected local coherence amongst CINs, as the discharge of pCINs in DMS did not coincide with ACh transients in DLS during immobility, despite showing similar pairwise synchrony within DMS compared to pCINs within DLS (**Extended Data Fig. 8**). Together, these data show that coherent CIN firing drives periodic ACh fluctuations, the amplitude of which reflects the degree of local spike coherence amongst neighboring CINs.

### Glutamatergic inputs drive periodic ACh fluctuations

How does this coherence arise? Striatal CIN receive convergent excitatory afferents from distributed cortical and thalamic brain regions^43^; we therefore postulated that synchronized glutamatergic inputs may coordinate CINs to produce periodic fluctuations in extracellular ACh levels. To test this, we imaged ACh while recording striatal spiking activity and local field potential (LFP) oscillations, which largely reflect the activity of afferent inputs^44-46^. ACh fluorescence was strongly modulated with respect to LFP oscillations in the 0.5–4 Hz frequency band (**Fig. 4m**), suggesting that coherence amongst CINs may be imparted by striatal afferents. Indeed, the vast majority of pCINs recorded (94.6%) were significantly entrained to 0.5–4 Hz LFP oscillations with a preferred phase of 218 ± 6 degrees (**Fig. 4n, o**), consistent with negative LFP deflections reflecting an increase in excitatory drive to pCINs. To further test whether glutamatergic inputs drive periodic ACh fluctuations, we delivered a cocktail of ionotropic glutamate receptor (iGluR) blockers locally in DLS while imaging ACh3.0 (**Fig. 4p**). This treatment did not significantly decrease net ACh fluorescence compared to saline, (**Extended Data Fig. 5o**), in line with the fact that CINs continue to fire cell-autonomously in vivo in the absence of glutamatergic drive^47^. By contrast, iGluR blockers rapidly and reversibly abolished spontaneous as well as movement- and reward-evoked ACh transients (**Fig. 4q-s**). Collectively, these data reveal that periodic fluctuations in ACh are driven by extra-striatal glutamatergic afferents, which act to synchronize the activity of CINs in striatum.

## Discussion

It is commonly assumed that striatal circuits bathe in a steady concentration of DA and ACh contributed by the tonic, cell-autonomous pacemaking of midbrain DANs and striatal CINs^48,49^. This neuromodulatory ‘soup’ is occasionally interrupted by phasic increases or decreases in extracellular DA and ACh levels driven by sensory and motor stimuli that play important roles in reinforcement learning and motor performance^9,14,18,19,50,51^. Here, we show that these phasic signals are prevalent when mice are immobile and in the absence of salient stimuli, with striatal DA and ACh levels fluctuating periodically several times a second. This observation, which hinges on the recent development of fast and specific sensors for DA and ACh with signal-to-noise sufficient to resolve single-trial responses calls into question the concept of a stable basal neuromodulatory tone in striatum. Importantly, spontaneous changes in DA and ACh levels do not occur randomly: they are locally coherent at 0.5–4 Hz, with DA signals maintaining a 90-degree phase-shift relative to ACh across behavioral contexts. ACh is therefore unlikely to serve as a context-specific gate or coincidence signal that disambiguates DA transients meant to promote learning vs. movement^12,13^. In addition, this finding suggests that rewards and other salient stimuli do not create unique neuromodulatory dynamics in striatum, but instead engage a common existing neural architecture that intrinsically supports rhythms in DA and ACh.

Surprisingly, the rapid, sub-second coordination between DA and ACh does not depend on their intra-striatal interactions, despite strong evidence in brain slices by us and many others^24-28^. It is possible that CINs in vivo do not readily drive action potentials in DA axons given their sustained firing, prominent synaptic depression, and propensity for nAChRs to desensitize. Another possibility is that DA and ACh release in vivo is not as synchronous as the electrical and optogenetic stimulations required to evoke DA release ex vivo^25-27^. That said, we cannot rule out that intra-striatal interactions between DA and ACh play an important role in other behaviors, on subcellular scales not resolved by photometry, or on slower time courses than those investigated here.

Our data support a model where spontaneous rhythms in striatal DA and ACh are inherited from extra-striatal afferents. Glutamatergic inputs are sufficient to drive phasic elevations in ACh^52^ and may contribute to pauses in CIN firing via feed-forward inhibition^53^ or periodic withdrawal of excitatory inputs^54^. Importantly, the magnitude of ACh fluctuations correlates with the degree of population synchrony amongst CINs, which increases when the brain produces concerted movements^23^ or responds to salient sensory stimuli^8^. Striatal ACh levels may therefore be construed as a moment-to-moment measure of coherence in the activity of distributed cortico- and thalamo-striatal afferents. Midbrain DANs similarly integrate excitatory (and inhibitory) inputs from many brain areas^42^ and exhibit significant baseline correlations in spiking that increase in strength upon reward presentation and learning^37^ and account for phasic changes in striatal DA levels more so than bursting of individual neurons^40,41^. Thus, DA and ACh both reflect the concerted dynamics of widely distributed neural circuits^55-57^, providing a novel framework for understanding how the basal ganglia motivate and reinforce behavior.

First, the presence of spontaneous reward-like rhythms in DA and ACh suggests that the established functions of both modulators in learning and synaptic plasticity extend beyond the time of reward delivery. Indeed, learning and actions are not exclusively driven by external rewards and salient sensory stimuli. Second, the phase offset between DA and ACh may create temporally-defined windows for learning; synaptic plasticity may preferentially occur when direct- and indirect-pathway SPNs fire within specific phases of this endogenous rhythm^17,58^, with repeated reactivation of SPN ensembles during post-learning rest offering a potential solution to the credit assignment problem^59,60^. Third, intrinsic fluctuations in DA and ACh may help specify the timing and vigor of self-paced volitional actions^10,18,19,23^ as well as sustain cognitive processes that lack overt behavioral correlates^61-63^. Thus, the autonomously-coordinated rhythms in DA and ACh revealed here provide a lens through which to understand the structure of activity within the basal ganglia as well as its impact on behavior in health and in neuropsychiatric disorders.

## Methods

### Animals

All procedures were performed in accordance with protocols approved by the NYU Langone Health (NYULH) Institutional Animal Care and Use Committee (protocol #170123). Mice were housed in group before surgery and singly after surgery under a reverse 12-hour light-dark cycle (dark from 6 a.m. to 6 p.m.) with *ad libitum* access to food and water, except when water-restricted to incite consumption of water rewards. Experiments were carried out using both male and female mice at 8–24 weeks of age. For conditional deletion of choline acetyl-transferase (ChAT) from forebrain cholinergic interneurons (CINs), a floxed ChAT line was crossed with Nkx2.1-Cre transgenic mice (both kindly provided by Dr. Robert Machold, NYULH), as described previously^64^. For conditional deletion of the β2 nAChR subunit from dopamine (DA) neurons, a floxed β2 line^65^ (kindly provided by Dr. Michael Crair, Yale) was crossed with a DAT-ires-Cre knockin mice^66^ (Jackson Laboratory strain #006660). C57BL6/J mice were used as control animals (Jackson Laboratory strain #000664).

### Stereotaxic surgery

Mice were anaesthetized with isoflurane, placed in a stereotaxic apparatus (Kopf Instruments) on a heating blanket (Harvard Apparatus) and administered Ketoprofen (10 mg/kg, subcutaneous). The scalp was shaved and cleaned with ethanol and iodine solutions before exposing the skull. For simultaneous recordings of DA and acetylcholine (ACh) in the dorsolateral striatum (DLS), craniotomies were drilled above the injection site (from bregma, in mm: AP +0.5, ML +2.5 for 0° angle implants or AP +0.5, ML +3.25 for 30° angle implants). 200nL of a 1:1 mix of AAV9-hSyn-ACh3.0 (ref. ^32^; titer 1.46 e+13; Vigene Biosciences) and AAV9-hSyn-rDA1m (ref. ^31^; titer 1.91 e+13; Vigene Biosciences) was injected 2.0mm below dura (0° angle) or 1.6mm below dura (30° angle) at a rate of 100 nL/min using a microsyringe pump (KD Scientific; Legato 111) fitted with a Hamilton syringe (1701N, gastight 10 μL) connected to a pulled glass injection micropipette (100 μm tip; Drummond Wiretrol II) via PE tubing filled with mineral oil. Injection micropipettes were left in place for 5 min before removal. The optimal dilution of each AAV was empirically determined to enable strong expression while avoiding toxicity. A custom-made 400 μm diameter optic fiber (Thorlabs FP400URT, NA = 0.5) connected to a ceramic ferrule (Thorlabs CFLC440) was implanted above the injection site to and cemented to the skull using C&B metabond (Parkell). Only fibers with 75 to 95% transmission efficiency were selected for surgical implantation. For mice used for simultaneous recording of ACh and DA in DLS and ipsilateral DMS, additional craniotomies were drilled above the DMS injection site (from bregma, in mm: AP +0.7, ML +1.3 for -5° angle implants). 200nL of a 1:1 mix of AAV9-hSyn-ACh3.0 and AAV9-hSyn-rDA1m was injected 2.0mm below dura and a second optic fiber was implanted at a depth of 1.85mm below dura. For mice used for acute intra-striatal pharmacological infusions in DLS, a custom guide cannula (Plastics One; 26 gauge) was chronically implanted ipsilateral to the chronic optic fiber implant at AP +0.5 and ML +1.7 at a - 10° angle at a depth of 1.75mm below the pial surface. The guide cannula was cemented to the skull using C&B metabond and covered with a dummy cannula. All mice had a custom titanium headpost implanted over lambda using C&B metabond to allow head fixation during recordings. Mice were allowed to recover in their cage for two weeks before head-fixation habituation, treadmill and lick spout training, and recordings.

For mice used for acute electrophysiological recordings in DLS, a second surgery was performed after the post-recover period, head-fixation habituation, and treadmill training. Mice were head-fixed using the previously cemented titanium headpost. A craniotomy centered on ML +2.0mm and extending from AP -0.5mm to +1.5mm was drilled and dura was carefully removed 24 h prior to recording. For acute electrophysiological recordings in dorsomedial striatum (DMS), the craniotomy was centered on ML +1.0mm. A second craniotomy (ML -1.0 mm, AP -1.0 mm) was drilled for grounding during recordings. Both craniotomies were covered with silicone sealant (Kwik-Cast) until recording. After craniotomies, mice were allowed to recover in their cage for one day before recordings.

For experiments evaluating the effects of DA neuron loss on entrainment of ACh release, an additional stereotaxic surgery was performed under isoflurane anesthesia after baseline imaging sessions. Desipramine (25 mg/kg) and pargyline (5 mg/kg) were administered intraperitoneally prior to surgery to increase the selectivity and efficacy of 6-hydroxydopamine (6OHDA) lesions^67^. A small craniotomy was performed above the substantia nigra pars compacta ipsilateral to the imaged striatum (−3.1 mm posterior from bregma, 1.3 mm lateral, marked during first stereotaxic surgery) and 3 μg of 6-OHDA (total volume: 200 nl) was injected

4.2 mm below dura at a rate of 100 nl/min. Mice were allowed to recover in their cage for two weeks, with twice daily intraperitoneally injections of glucose (5% in 1mL 0.9% saline), twice daily subcutaneous injections of 0.9% saline (1 mL), and once daily subcutaneous injections of ketoprofen (10mg/kg in 0.9% saline).

### Behavior Analysis

Mice were habituated to head-fixation in the rig in a dark soundproof chamber and locomotion on the cylindrical wheel for a minimum of 5 days prior to any recording. A subset of mice had access to water within their home cage restricted to incite consumption of water delivered from a spout at semi-random intervals during imaging sessions. Licking was monitored using a capacitive touch sensor (Sparkfun AT42QT1010). Solenoid valve (Lee Company LHQA0531220H) opening and lick sensor signals were acquired as digital inputs to a National Instruments acquisition board (NI BNC-2090A) at a sampling rate matching the photometry signal acquisition rate using Wavesurfer software. Reward delivery and lick event times were recorded as time points exceeding a threshold of 0.5 V. Analyses of reward-evoked DA and ACh transients were limited to events where mice withheld licking for at minimum 0.5 s prior to solenoid opening and then collected water within 250 ms. On average, the first lick occurred within 164 ± 12 ms of solenoid opening. Reward period was defined as the 1 s period following solenoid valve opening.

Treadmill velocity was extracted from positional information provided by a rotary encoder (MA3 magnetic shaft encoder, US Digital), down-sampled to 50 Hz for all recordings. Immobility was defined as any period of time lasting at least 4 s during which velocity does not exceed 0.25 cm/s beginning at minimum 0.5 s after treadmill velocity decreases below 0.25 cm/s and ending 1 s before treadmill velocity exceeds 0.25 cm/s again. Solenoid openings, licking responses and postural adjustments on the wheel (e.g. grooming, detected using an infrared camera (Sony AKK CA20 2.8mm FPV camera) were excluded from periods of immobility. Movement bouts were defined as any period of time when absolute treadmill velocity exceeded 4 cm/s for a minimum of 4 s, preceded and followed by 4 s of immobility. Treadmill acceleration was calculated as the first derivative of smoothed (*movmean* function in MATLAB; window, 0.2 s) treadmill velocity. Peaks of positive acceleration were determined by finding the time points of local maxima (*findpeaks* function in MATLAB with optimized thresholds, minimal prominence: 0.5 cm/s^2^) during periods of active locomotion.

### Fiber Photometry Recording

To excite the ACh3.0 and rDA1m sensors, we used fiber-coupled LEDs at 470 nm (Thorlabs M470F3) and 565 nm (Thorlabs M565F3), respectively. Excitation light was passed through a fiberoptic patch cord (Doric 400 μm 0.48 NA) to a fluorescence mini-cube (Doric FMC5_E1(460-490)_F1(500-540)_E2(555-570)_F2(580-680)_S) and connected to the chronic optic fiber implant in the mouse via a fiberoptic patch cord (Doric 400 μm 0.48 NA). Emission light was collected through the same patch cord and fluorescence mini-cube connected to two photoreceivers (Newport 2151) using a fiberoptic patch cord (Doric 600 μm 0.48 NA); one for green emitted light, and another for red emitted light. Excitation light was delivered in one of two modes: continuous wave (CW) or frequency modulated (FM). We used the former to image brains with a single sensor expressed and when performing concurrent *in vivo* extracellular recordings to minimize optoelectric artefacts. FM mode was used for imaging DA and ACh sensors simultaneously to minimize cross-talk between fluorescence channels. In CW mode, light power measured (Thorlabs PM100D) at the tip of fiber optic patch cord was set to 20-60 μW by manually adjusting the LED driver before each recording. In FM mode, a sinusoid generated using Wavesurfer software (Janelia) and outputted from a National Instruments acquisition board (NI BNC-2090A) was used to drive the fiber-coupled LED. The 473 nm and 565 nm LEDs were modulated at either 217 Hz or 319 Hz, alternating between recording sessions. Light power was set by adjusting amplitude and offset parameters of the sinusoidal control voltage by monitoring the output voltage from the photoreceiver using Wavesurfer software to achieve a sinusoid peak-to-peak amplitude of 1-2V. After each recording session, light power (18-62 μW) was measured at the tip of the fiberoptic patch cord. We ensured that imaging in FM mode does not generate artefactual oscillations by repeating key observations in mice with only one fluorescent sensor expressed at a time in CW mode. Photometry signals read out by the photoreceiver were digitized at 2 kHz (for CW mode) or 5 kHz (for FM mode) by a National Instruments acquisition board (NI BNC-2090A) or at 30 kHz (for combined fiber photometry and acute electrophysiology recordings) by an Open Ephys acquisition board.

### Fiber Photometry Analysis

#### Photometry signal processing

Raw photometry signals collected as a voltage from the photoreceiver were processed using custom MATLAB code. For recordings in CW mode, raw voltage signals were first low-pass filtered at 20 Hz using a butterworth filter to remove high-frequency noise. The filtered signal was then down sampled to 50 Hz (above the Nyquist frequency to prevent aliasing), and the final photometry signal (outputted as a % value) was obtained using the equation Δ*F/F = (F – F*_*0*_*)/F*_*0*_, in which the *F*_*0*_ is baseline fluorescence. The latter was computed by interpolating the bottom percentile of fluorescence values measured in 10 s-long sliding windows (0% overlap) along the entire photometry trace. For recordings acquired in FM mode, a copy of each sinusoid generated in Wavesurfer was recorded as a reference signal alongside the modulated green and red fluorescence signals from the photoreceivers. We demodulated fluorescence signals as follows: First, each fluorescence signal was band-pass filtered (reference sinusoid frequency ± 10 Hz; Butterworth; order = 6) to isolate the main frequency component. If the reference and fluorescence signals were in-phase, a phase dependent modulation was performed by multiplying the reference and modulated signal^68^, which would remove the frequencies and phases of the two signals. More commonly, the reference signal and fluorescence signal had similar frequency components but were slightly phase-shifted. In this case, we used a standard quadrature demodulation method to demodulate the fluorescence signal^69^. The demodulated fluorescence signal was then down sampled to 50 Hz and baseline adjusted as described above.

#### Frequency analysis

Analysis of frequency components enriched in the ACh and DA photometry signals was performed on raw, unfiltered, non-demodulated, non-downsampled fluorescence signals from time periods when mice were either immobile or locomoting. We computed the fast Fourier transform of these data using the *fft* function in MATLAB and extracted the single-sided amplitude (power) spectrum across frequency domains spanning 0 to 0.5*acquisition sampling frequency. Power spectra shown in Fig. 1i, m were normalized by subtracting the power at 100 Hz and dividing by the power at 0.1 Hz. To isolate power specifically contributed by the ACh3.0 and rDA1m sensors, we then subtracted the mean normalized power spectra of the inert fluorophore GFP or tdTomato, respectively, across the same frequency domains, as shown in Fig. 1j, n. To statistically compare ACh3.0 and rDA1m power across mice and conditions, we computed the integral of the power spectrum within the 0.5-4Hz frequency band, as shown in Fig. 1k, o.

#### Cross-correlation and coherence analysis

To correlate ACh3.0 and rDA1m fluorescence, we calculated the Pearson correlation coefficient *r* between the two photometry signals using the *xcorr* function in MATLAB (window, 1 s; bin, 20 ms) after parsing them based on behavioral state (i.e. reward, locomotion, or immobility). Confidence intervals were computed by repeatedly calculating Pearson’s *r* after one of the photometry signals was shifted in time (increment, 1 s; repeats, 100) and then extracting the 5th and 95th percentiles across the correlation window for each bin. The correlation coefficient *r* and lag at the maximal absolute deflection were computed and compared across behavioral states and/or cohorts, as shown in Fig. 2b, c. To quantify how closely DA and ACh fluorescence signals co-varied in time and the relative timing between the two signals we computed their coherence and extracted the magnitude of the coherence and phase offset, respectively. We performed this over different frequency domains to examine in which frequency band coherence is greatest, as well as over time domains to examine whether the degree of coherence and phase offset between DA and ACh varied between behavioral states. We computed coherence using a custom MATLAB script adapted from code acquired from the Buzsaki lab (https://github.com/buzsakilab/buzcode). Briefly, we calculated the magnitude and phase of the DA-ACh coherogram using a multi-taper estimation with the chronux function *cohgramc* (window, 10 s; overlap, 5 s; step, 5; padding, 0). We computed confidence intervals by randomly shifting one of the photometry signals (increment, 1 s; repeats, 100) and extracting the 5th and 95th percentiles of the magnitude and phase of the resultant coherogram for each repetition. To compare coherence at different frequencies, we averaged coherence magnitude and phase offset across the time dimension, as shown in Fig. 2d, f. To compare each between behavioral states, we calculated the median coherence magnitude and phase offset within the 0.5-4 Hz frequency band, as shown in Fig. 2e, g. Phase offset was only examined if a significant effect of coherence magnitude was observed.

#### Phase analysis

To estimate instantaneous phase across time, each photometry signal was filtered (0.5-4 Hz; Butterworth; order, 3) and the phase angle of the Hilbert transform was extracted using the *hilbert* function in MATLAB. We next extracted the amplitude of the DA and ACh fluorescence signals corresponding to each oscillatory cycle (from -180 to +180 degrees; bin size, 10 deg.) and calculated the mean amplitude within each bin. To highlight the temporal relationship between DA and ACh across behavioral states, we normalized fluorescence amplitude between 0 to 1 across each full cycle, as shown in Fig. 2j. We applied the same process to determine the relationship between the phase of ACh fluctuations and wheel acceleration, as shown in Extended Data Fig. 6c.

### Acute Electrophysiological Recording

Electrophysiological recordings were made with silicon microprobes^70^ (128D; IDAX microelectronics) affixed to metal rods acutely lowered into the brain using a micromanipulator (Scientifica PatchStar). For DLS recordings, probes were inserted perpendicularly to brain surface at approximately AP +0.0 to +1.0mm and ML +2.0 or +2.5mm to a depth of about 2.5-3.0 mm. Prior to insertion on each recording day, the electrode shafts were coated with fluorescent dye (DiI, Thermo Fisher Scientific V22885) for *post-hoc* identification of probe insertion location. Probes were lowered at a speed of 1 µm/s to limit tissue damage and, once at the desired depth, were left in place for at least 45 min prior to initiating a recording to allow for the tissue to settle and minimize unit drift. Electrophysiological data were recorded with Open Ephys at a sampling rate of 30 kHz. Wheel positional encoder and, when applicable, photometry data were also recorded using an I/O board connected to the Open Ephys acquisition board at a sampling rate of 30 kHz.

### Acute Electrophysiological Analysis

#### Spike sorting and putative cell type determination

Raw electrophysiological data was spike-sorted using Kilosort 2 (www.github.com/MouseLand/Kilosort2) and the resulting spike clusters were visualized and manually curated using Phy2 (https://phy.readthedocs.io/en/latest). High-quality single units were identified based on the following inclusion criteria: (1) waveform trough to peak amplitude exceeding 80 μV; (2) a minimum of 500 spikes recorded per 5400 s recording session; (3) waveform peaks preceding waveform troughs; (4) presence of a clear refractory period, assessed by considering the length of time that the firing rate of a unit is suppressed following a spike in auto-correlograms (window, 1 s; bin, 1 ms). Units passing these quality criteria were then classified into three putative striatal cell types: putative striatal projection neurons (pSPNs), putative cholinergic interneurons (pCINs), and a third class of unidentified neurons (others, which likely includes fast-spiking interneurons and other GABAergic interneurons) using established criteria^71-76^ such as firing rate, coefficient of variation, phasic activity index, and spike waveform duration. The firing rate of a unit was calculated as the number of spikes divided by the duration of the recording. The coefficient of variation (CV) was defined as the standard deviation of the distribution of inter-spike intervals (ISIs) divided by the mean of the distribution. The index of phasic activity was calculated as the fraction of recording time containing ISIs longer than two seconds. The waveform duration was calculated as the time between the maximum downward spike deflection to the following upward peak. For analyses during immobility or locomotion, we extracted spikes that occurred during windows of time when the animal was determined to be immobile or actively locomoting, respectively, using criteria described in Behavior Analysis above.

#### Spike train cross-correlation

In recordings with ≥ 2 simultaneously recorded pCINs, we first generated pairs using the *nchoosek* function in MATLAB. For each pair of pCINs, one was randomly selected to be the reference unit and we calculated the spike cross-correlogram (CCG) for that pair as a count of the number of spikes emitted by one pCIN relative to the spikes of the reference unit (window, 2 s; bin, 10 ms). We normalized spike counts to a firing rate by dividing by number of spikes within each time bin by the total number of reference unit spikes and by bin size: firing rate = (spike counts/N^ref^)/bin duration (s). To allow comparisons across units with different baseline firing rates, we normalized instantaneous firing rates to background, baseline rates by subtracting the mean firing rate of the non-reference unit during a given behavioral state before dividing the resultant value by the same mean: Normalized instantaneous rate = (instantaneous rate – µ)/µ. To compute confidence intervals, we ran a CCG using the same reference unit and a shuffled spike train of the second pCIN. Each shuffled spike train was generated following random permutation of a unit’s ISIs and the 2.5th and 97.5th percentiles were extracted across CCGs. Lastly, to assess the proportion of units that significantly increase their firing rate, we identified all the time bins at various lags where the CCG output was either above or below the 95% confidence interval.

#### Firing rate change at event times

For recordings where ACh fluorescence was imaged concurrently with spiking activity, we determined the instantaneous firing rate of pCINs relative to peaks and troughs in ACh by filtering photometry signals (0.5-4 Hz; Butterworth; order = 3), identifying transients larger than 2 standard deviations from the filtered signal, and computing a peri-event time histogram by aligning spike times to event times such as ACh peaks and ACh troughs. Spike times were counted in 20 ms bins (determined based on the final sampling rate of the photometry signal, 50 Hz) across a two second time window centered on the event times, and then averaged across all events to generate a rate. We normalized firing rate change at event times to the mean firing rate for a given behavioral state to allow for comparison between units of different firing rates. To obtain confidence intervals, a shuffled spike train was instead aligned to event times. To assess the proportion of units that significantly change their firing rate, we determined the number of bins where a unit’s firing rate either exceeds or falls below the 95% confidence interval.

#### Coherent spiking analysis

Only recordings with ≥ 2 pCINs and concurrent imaging of ACh fluorescence were included. Spike trains were first parsed to include only spikes occurring during immobility, as defined in Behavioral Analysis. For each spike of a selected reference pCIN, we determined if other pCINs discharged within ± 10 ms by generating peri-event counts relative to spikes. The resulting output was a matrix of binary values, with the number of columns reflecting the length of the spike train of the reference unit and the number of rows corresponding to the number of concurrently recorded units (N). This output matrix was then summed across all rows, such that the range of values of the final vector would be 1 to N. Lastly, spike times of the reference unit were stratified based on the value within each bin. Bins where only 1 out of N units spiked reflect low synchrony amongst the population of pCINs, whereas bins with a value of N out of N possible units corresponded to synchronous activity across all concurrently recorded pCINs.

Simultaneously recorded ACh photometry signal was aligned to stratified spike times of the reference unit, and a spike-triggered average was computed by averaging the photometry signal across each group (1/N to N/N), as shown in Fig. 4k. To compare units across multiple recordings with a variable total number of pCINs, only 1/N and N/N spike-triggered averages were used, as shown in Fig. 4l. For computation of confidence intervals, the simultaneously recorded photometry signal was aligned to spike times in a shuffled spike train.

#### LFP analysis

Local field potential (LFP) signals were extracted from wideband electrophysiological recordings from one striatal channel and down-sampled to 1250 Hz. The LFP was then filtered at 0.5-4 Hz and the instantaneous phase was determined using a Hilbert transform. Phase locking was determined by calculating the instantaneous phase angle at each spike time for a given unit during immobility. A histogram of phase angles was calculated, and the circular mean and resultant vector were calculated for each unit, as shown in Fig. 4n. Only units that were significantly modulated (Rayleigh test) were included in the analysis. For recordings where ACh fluorescence was imaged concurrently, phase locking was determined by calculating a histogram of photometry signal amplitudes at each phase angle, and rescaling across each cycle, as shown in Fig. 4m. Cross-correlation between the filtered LFP signal and concurrently recorded photometry signal and was calculated as above.

### Immunohistochemistry

Mice were deeply anesthetized with isoflurane and perfused transcardially with 4 % paraformaldehyde in 0.1 M sodium phosphate buffer. Brains were post-fixed for 1–3 days and sectioned coronally (50–100 μm in thickness) using a vibratome (Leica; VT1000S). Brain sections were mounted on superfrost slides and coverslipped with ProLong antifade reagent with DAPI (Molecular Probes). ACh3.0, rDA1m, GFP, and/or tdTomato fluorescence were not immuno-enhanced. Whole sections were imaged with an Olympus VS120 slide scanning microscope, as shown in Extended Data Fig. 1a.

### Reagents

Drugs (all from Tocris) were reconstituted and stored according to the manufacturers’ recommendations. For systemic antagonism of mAChRs and D2Rs (Fig. 1h-o), scopolamine (2 mg/kg) and sulpiride (30 mg/kg) were respectively prepared daily in sterile physiological saline (0.9% NaCl) and administered intraperitoneally 30 min prior to the start of photometry recordings. For local injections within DLS, an internal cannula (Plastic One; 33-gauge) was connected to a syringe (Hamilton; 1701N, gastight 10 μL) through a tube containing either sterile saline (RECIPE) or a pharmacological agent dissolved in saline and was inserted into the guide cannula before the recording session. Drug infusion controlled by a microsyringe pump (KD Scientific; Legato 111; rate: 100nL/min) was initiated 10-20 min after baseline recording. The following drugs were used: 1 mM NBQX (2,3-Dioxo-6-nitro-1,2,3,4-tetrahydrobenzo[*f*]quinoxaline-7-sulfonamide) + 5 mM D-AP5 (D-(-)-2-Amino-5-phosphonopentanoic acid) to block iGluRs, 400 μM SCH23390 to block D1Rs, 600 μM sulpiride to block D2Rs, 5 mM DHβE (dihydro-β-erythroidine hydrobromide) to block nAChRs and 20 mM scopolamine to block mAChRs.

### Statistics

Data are reported in text and figures as mean ± standard error of mean (sem), with shaded areas and error bars in figures representing sem. Data were compared using MATLAB with the following statistical tests (as indicated in the text): Student’s paired *t*-test for comparisons between paired data points, Student’s two-sample *t*-test for comparisons between un-paired data points, and one-way analysis of variance (ANOVA) followed by Dunn’s Multiple Comparison Test for comparisons between multiple groups. For fiber photometry experiments, n-values represent the number of mice. All imaging sessions obtained per condition from the same mouse (average: 2 sessions per condition; range: 1–4) were concatenated and analyzed as n = 1. For electrophysiology recordings, n represents the number of units or unit pairs. Exact p-values are provided in text and figure legends, and statistical significance in figures is presented as * p < 0.05, ** p < 0.01 and *** p < 0.001.

## Acknowledgments

We thank G. Buzsaki, M. Long, S. Shoham, R. Tsien and members of the Tritsch laboratory for comments on the manuscript, M. Crair (Yale) and R. Machold (NYULH) for providing the β2^f/f^ and ChAT^f/f^:Nkx2.1-Cre mice, respectively. We acknowledge the New York University Langone Health Rodent Genetic Engineering Laboratory for rederivation, the Genotyping Core Laboratory for mouse genotyping, the Department of Comparative Medicine for animal care and maintenance and the Neuroscience Institute’s imaging facilities for microscope availability. This work was supported by the National Institutes of Health (DP2NS105553 to NXT; T32NS086750, T32GM007308 and T32GM136573 to ACK), the Alfred P. Sloan, Dana, Whitehall and Feldstein Medical Foundations (NXT) and a Vilcek Scholars Award (ACK).

## Author contributions

ACK and NXT conceived of the project, designed and performed experiments, analyzed and interpreted the data, and wrote the manuscript. PM helped with analyses and YL provided reagents.

## Competing interests

YL is listed as an inventor on a patent application (PCT/CN2018/107533) describing GRAB probes. The other authors declare no competing interests.

Supplementary Information is available for this paper

All data, code, and materials will be made available upon publication. Correspondence and requests for materials should be addressed to NXT.

## Extended Data

**Extended Data Fig. 1.**
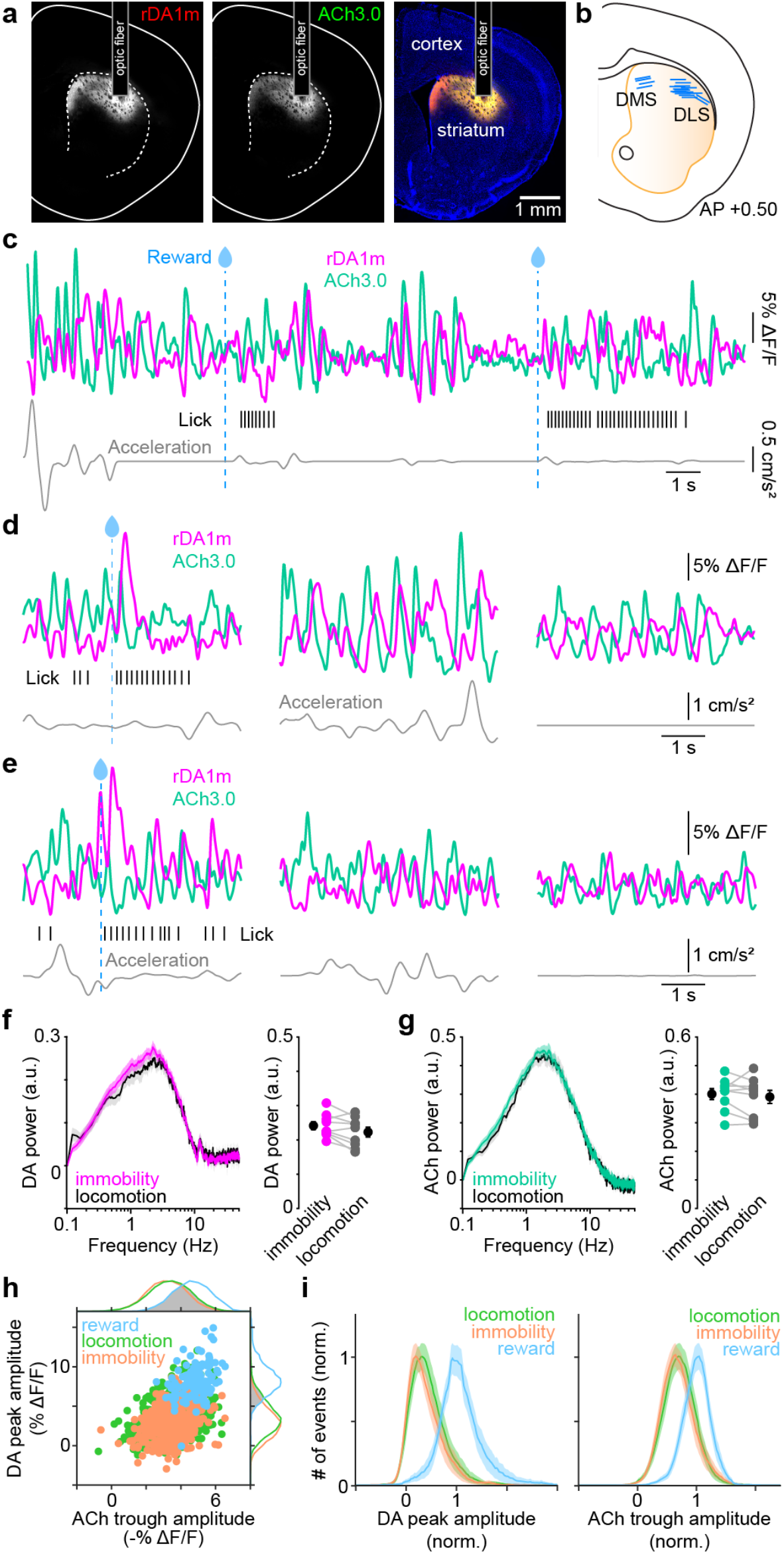
DA and ACh levels undergo spontaneous periodic fluctuations during reward, locomotion, and immobility. **a**, Epifluorescence images of rDA1m (*left*) and ACh3.0 (*middle*) expression in coronal section of striatum. *Right*, merged image overlaid with DAPI nuclear stain (blue). **b**, Schematic of striatum depicting tip of all recovered optic fiber implants in DLS and DMS. **c**, Example continuous recording of rDA1m (magenta) and ACh3.0 (teal) fluorescence (*top*), lick events (*middle*) and treadmill acceleration (*bottom*) from another mouse than that shown in Fig. 1. Water delivery highlighted with dashed blue line. **d**, Example reward (*left*), locomotion (*middle*) and immobility (*right*) photometry recording segments from same mouse as (**c**). **e**, Same as **d** for a different mouse. **f**, *Left*, power spectrum of DA signal during periods of immobility (magenta) and locomotion (black). *Right*, mean (± sem) DA power at 0.5-4 Hz (n = 13 mice; p = 0.05, Student’s paired *t*-test). **g**, Same as **f** for ACh (p = 0.5, Student’s paired *t*-test). **h**, Scatterplot of the amplitude of DA peaks and concurrent ACh troughs imaged during reward (blue), locomotion (green) and periods of immobility (orange) during one recording session. Amplitude distribution for ACh (*top*) and DA (*right*), with overlap across states shown in gray. **i**, Mean (± sem) distribution of DA peak (*left*) and ACh trough (*right*) amplitudes for all mice (n = 13) normalized to the mode of reward distributions showing considerable overlap between behavioral states.

**Extended Data Fig. 2.**
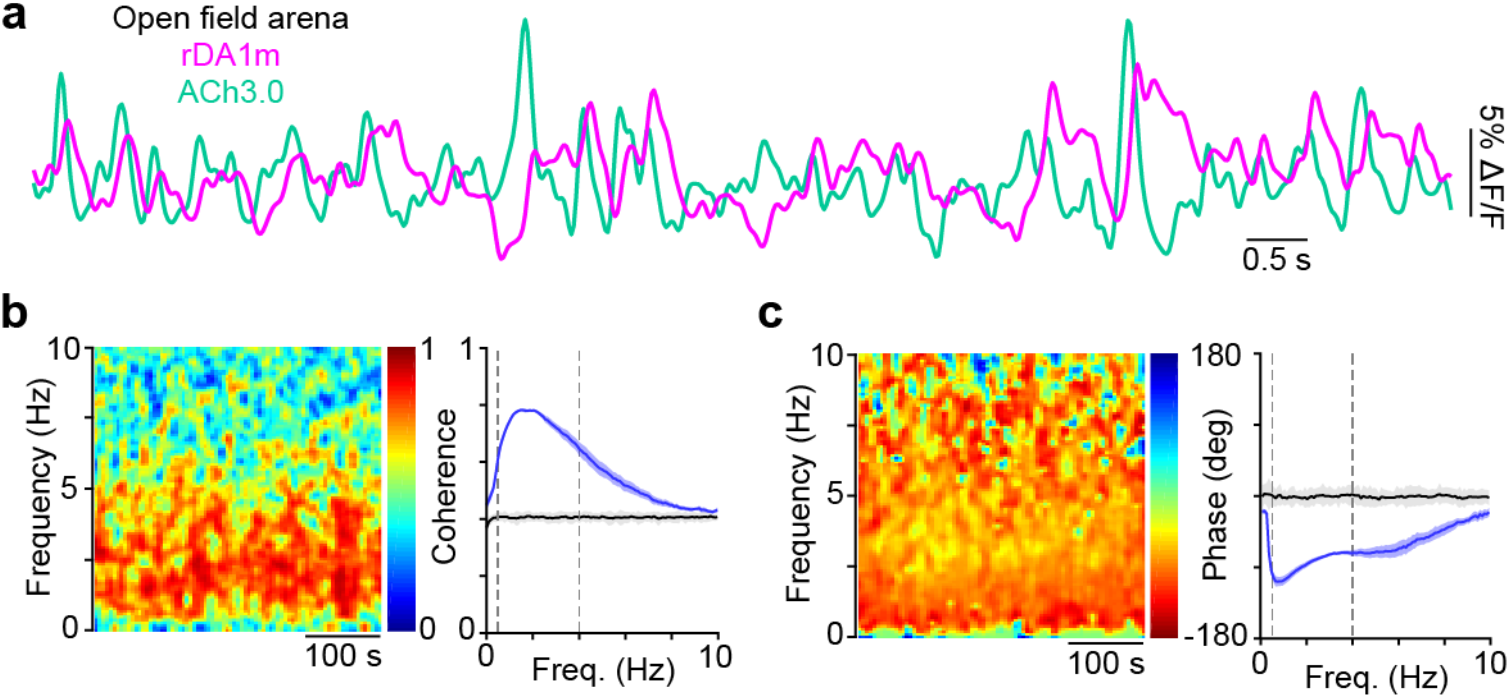
Spontaneous periodic fluctuations in striatal DA and ACh levels in unrestrained mice. **a**, Simultaneous photometry recording of DA (magenta) and ACh (teal) signals in a mouse freely behaving in an open field arena. **b**, *Left*, magnitude of coherence between DA and ACh signals across frequency and time domains during an open field recording. *Right*, population mean (± sem) coherence at different frequencies in n = 4 mice. Shuffled control shown in black. Vertical lines indicate 0.5 Hz, 4 Hz. **c**, Same as **b** for phase offset.

**Extended Data Fig. 3.**
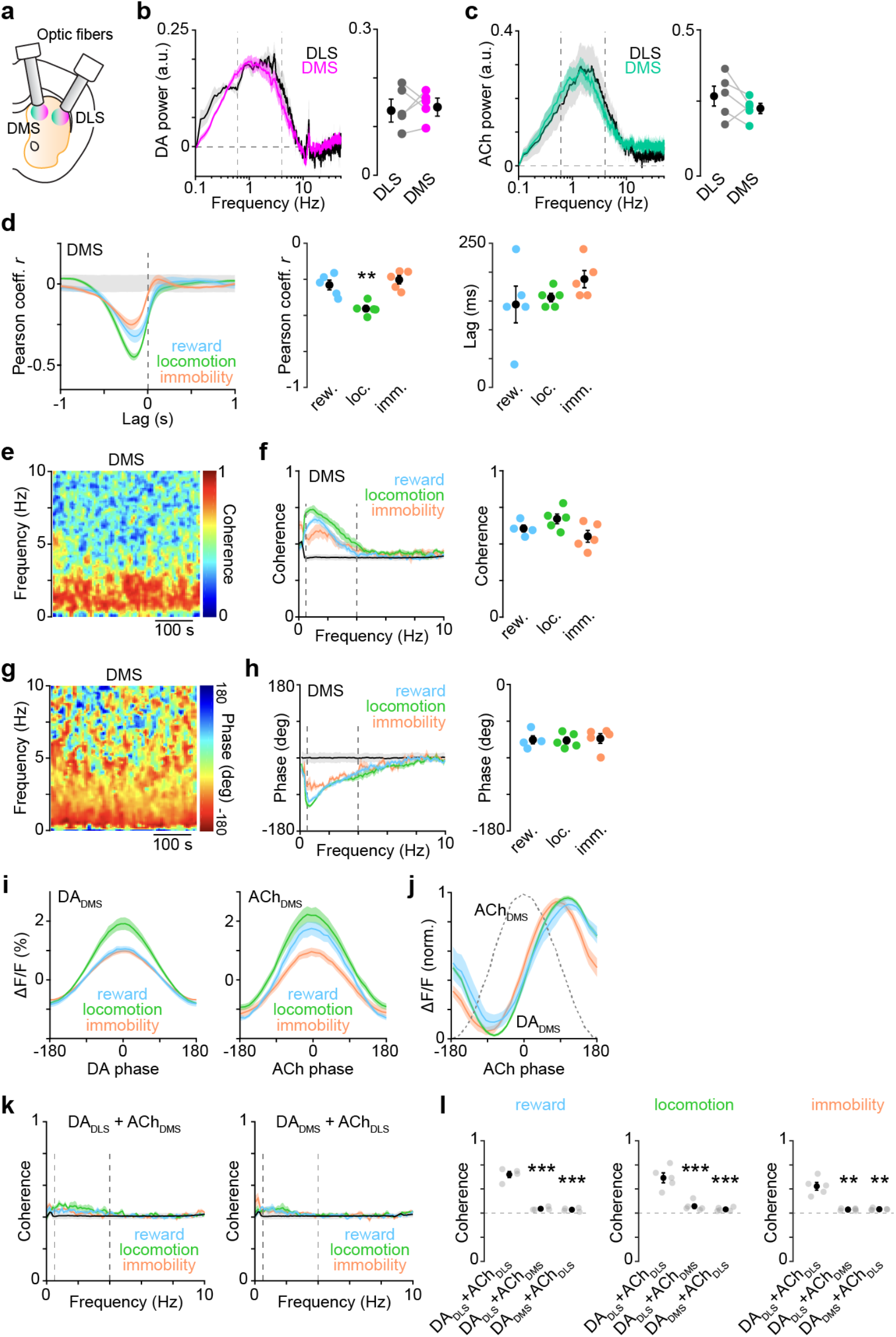
DA and ACh levels also fluctuate periodically in DMS, but are not coherent with DA and ACh fluctuations in DLS. **a**, Schematic of experimental preparation. **b**, *Left*, power spectrum of DA signal during immobility in n = 5 mice simultaneously recorded in DMS (DA_DMS_; magenta) and DLS (DA_DLS_; black). *Right*, mean (± sem) DA power at 0.5-4 Hz (p = 0.5, Student’s paired *t*-test). **c**, Same as **b**, for ACh in DMS (ACh_DMS_; teal) and DLS (ACh_DLS_; black) (p = 0.3, Student’s paired *t*-test). **d**, *Left*, mean (± sem) cross-correlation between simultaneously recorded DA_DMS_ and ACh_DMS_ signals during reward (blue), locomotion (green), and immobility (orange). *Middle*, peak correlation coefficient (Pearson’s *r*) in individual mice across behavioral states (**p = 0.006 vs. reward and immobility, Dunn’s multiple comparisons). *Right*, time lag of negative cross-correlation peak (p = 0.4, one-way ANOVA). **e**, Magnitude of coherence between DA_DMS_ and ACh_DMS_ signals across frequency and time domains for an example recording. **f**, *Left*, population mean (± sem) coherence at different frequencies during reward (blue), locomotion (green), immobility (orange). Vertical lines depict 0.5 and 4 Hz. *Right*, median coherence at 0.5-4 Hz band for individual mice across behavioral states. Population mean (± sem) shown in black (p = 0.09, one-way ANOVA). **g**, Same as (**e**), for phase offset between DA_DMS_ and ACh_DMS_. **h**, Same as **f**, for phase offset between DA_DMS_ and ACh_DMS_ (p = 0.2, one-way ANOVA). **i**, *Left*, mean (± sem) DA_DMS_ fluorescence at different phase of periodic DA_DMS_ fluctuations in the 0.5-4 Hz frequency band during reward (blue), locomotion (green), and immobility (orange). *Right*, same for ACh_DMS_ fluorescence vs. phase of ACh_DMS_ fluctuations. **j**, Peak-normalized DA_DMS_ fluorescence vs. phase of ACh_DMS_ fluctuations (shown as gray dotted line) in 0.5-4 Hz frequency band during reward (blue), locomotion (green), and immobility (orange). **k**, *Left*, mean (± sem) coherence between simultaneously recorded DA_DLS_ and ACh_DMS_ at different frequencies during reward (blue), locomotion (green), and immobility (orange). *Right*, same for coherence between DA_DMS_ and ACh_DLS_. **l**, Median coherence at 0.5-4 Hz band during reward (*left*), locomotion (*middle*), and immobility (*right*). Reward: DA_DLS_+ACh_DMS_ p = 8.14×10^−7^; DA_DMS_+ACh_DLS_ p = 6.07×10^−7^; locomotion: DA_DLS_+ACh_DMS_ p = 7.46×10^−5^; DA_DMS_+ACh_DLS_ p = 6.07×10^−5^; immobility: DA_DLS_+ACh_DMS_ p = 0.004; DA_DMS_+ACh_DLS_ p = 0.005; all vs. DA_DLS_+ACh_DLS_, Dunn’s multiple comparisons.

**Extended Data Fig. 4.**
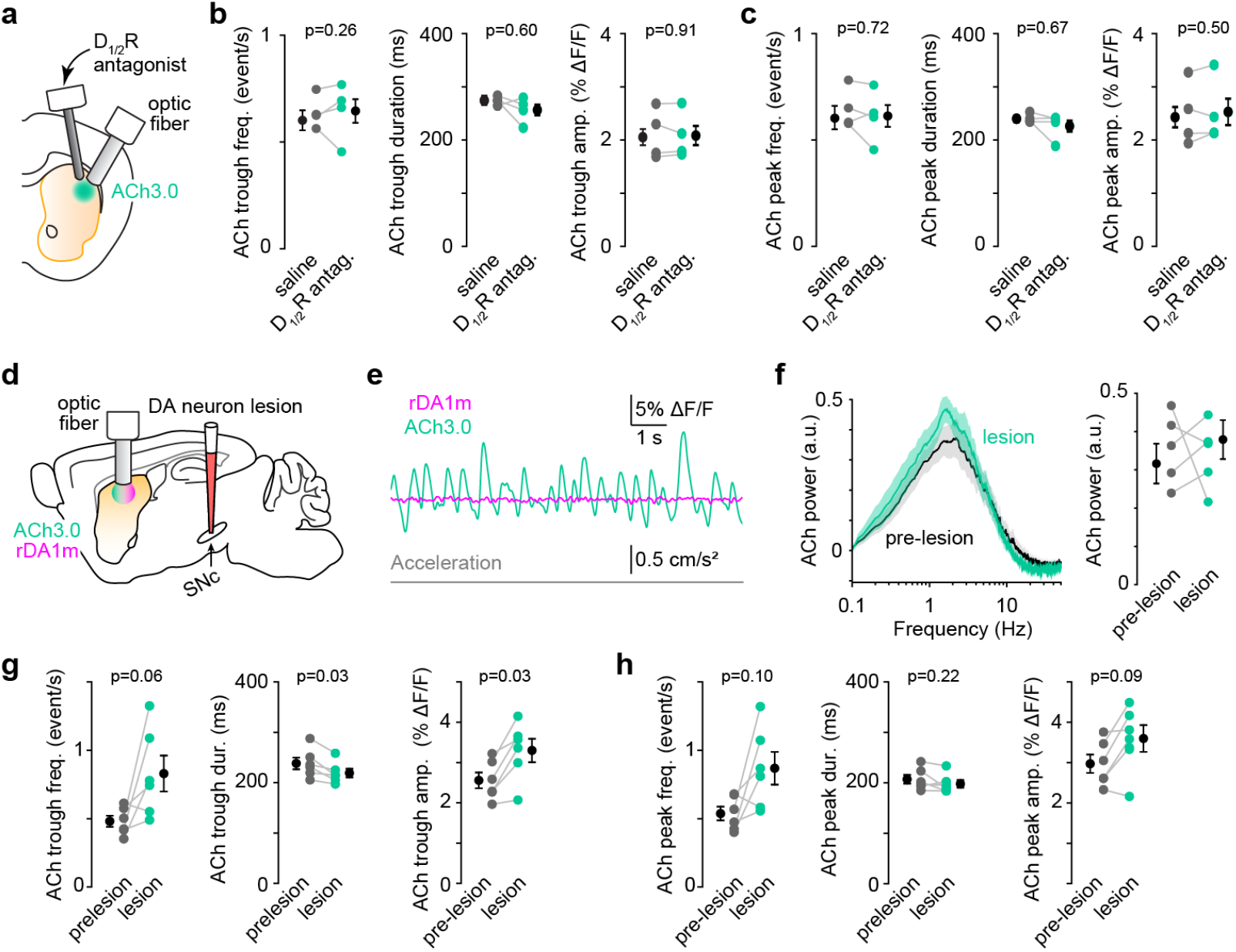
DA does not drive periodic fluctuations in ACh. **a**, Experimental preparation for local pharmacological inhibition of DA signaling in DLS while imaging ACh fluorescence. **b**, Frequency (*left*), duration (*middle*), and amplitude (*right*) of periodid dips in ACh fluorescence recorded during immobility after infusion of saline (gray) or D1/2R antagonists (teal). Population mean (± sem) shown in black (n = 4 mice). p-values (Student’s paired *t*-test) indicated in figure. **c**, Same as **b** for peaks in ACh fluorescence. **d**, Experimental preparation for 6OHDA-mediated lesioning of midbrain DA neurons while imaging DA and ACh fluorescence in DLS. **e**, Example DA (magenta) and ACh (teal) fluorescence traces following 6OHDA lesion during immobility (acceleration shown in gray). **f**, *Left*, power spectrum of ACh signal during immobility, before (gray) and after 6OHDA lesion (teal). *Right*, ACh power in 0.5-4 Hz frequency band in 5 mice. Population mean (± sem) shown in black (p = 0.44, Student’s paired *t*-test). **g**, Same as **b**, for periodic dips in ACh fluorescence before and after 6OHDA lesion, showing that chronic DA depletion causes phasic dips in ACh to become larger and slightly more frequent. This suggests a slow, negative influence of DA on ACh signaling, not a direct, sub-second role for DA in patterning ACh transients in DLS. **h**, Same as **b**, for peaks in ACh fluorescence before and after 6OHDA lesion.

**Extended Data Fig. 5.**
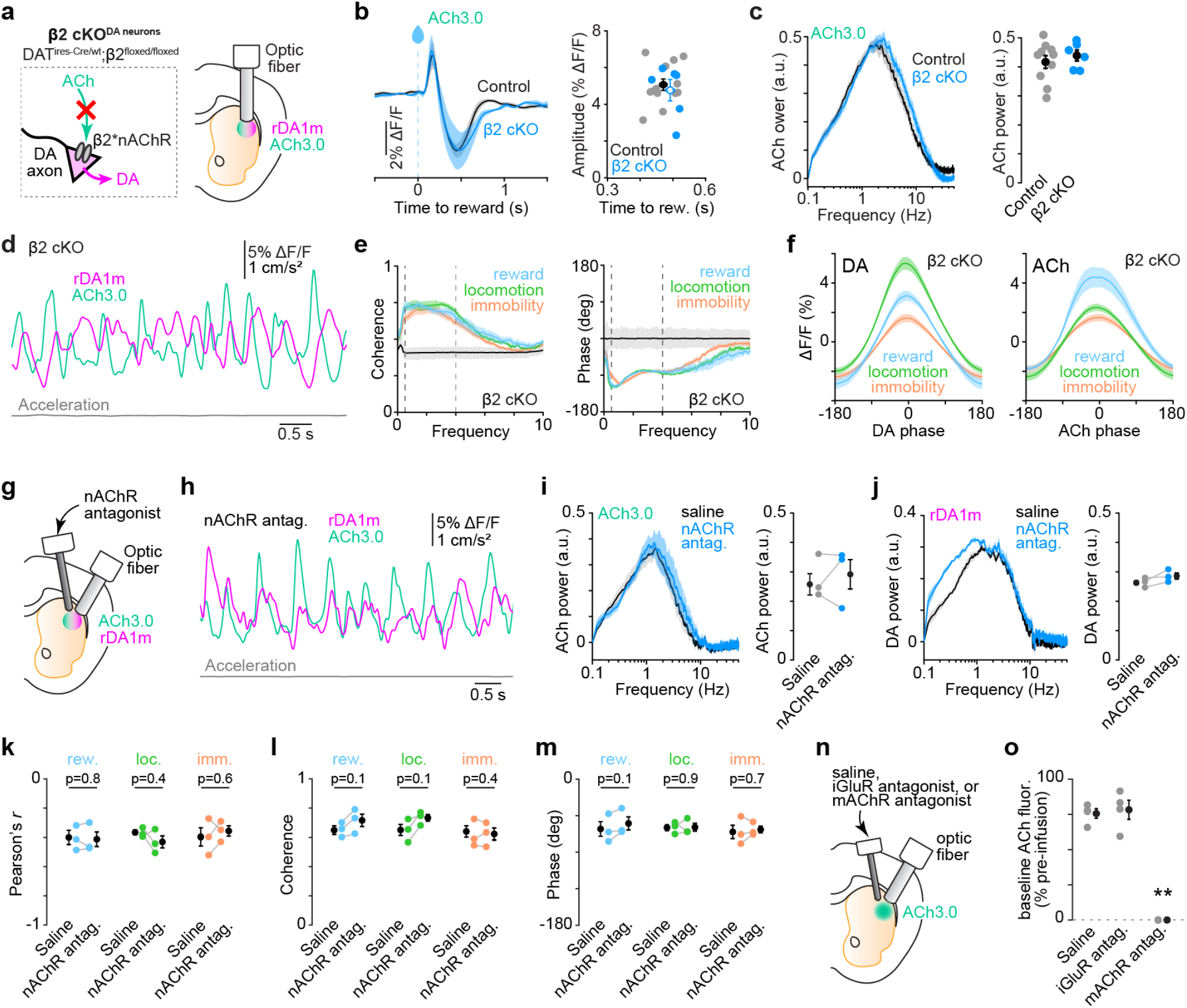
Nicotinic ACh signling does not drive periodic DA fluctuations. **a**, *Left*, conditional deletion of β2 nAChR subunit from DA neurons as a means to genetically prevent ACh-evoked DA release in striatum. *Right*, experimental preparation. **b**, *Left*, example reward-evoked ACh response imaged from DLS in a control (black) and a β2 cKO (blue) mouse. *Right*, scatter plot of reward-evoked ACh dip amplitude and timing in control (n = 13 mice; gray) and β2 cKO mice (n = 6 mice; blue). Group means (± sem) overlaid. Amplitude: p = 0.1; timing: p = 0.8; Student’s two-sample *t*-test. **c**, *Left*, power spectrum of ACh signal during immobility in control (black) and β2 cKO (blue) mice. *Right*, mean (± sem) ACh power in 0.5-4 Hz band; p = 0.5, Student’s two-sample *t*-test. **d**, Example DA (magenta) and ACh (teal) traces from a β2 cKO mouse, with acceleration shown in gray. **e**, *Left*, mean (± sem) coherence between DA and ACh signals in β2 cKO mice at different frequencies during reward (blue), locomotion (green), immobility (orange). Vertical lines depict 0.5 and 4 Hz. *Right*, same, for phase offset between DA and ACh. **f**, *Left*, mean (± sem) DA fluorescence at different phase of periodic DA fluctuations in the 0.5-4 Hz frequency band across behavioral states in β2 cKO mice (N = 6). *Right*, same for ACh fluorescence vs. phase of periodic ACh fluctuations. **g**, Experimental preparation for local pharmacological inhibition of nAChR signaling in DLS. **h**, Same as **d**, after DLS infusion of the nAChR antagonist DHbE. **i**, Same as **c** for ACh power after DLS infusion of saline vs. nAChR antagonist (n = 3; p = 0.8, Student’s paired *t-*test). **j**, Same as **i**, for DA power (p = 0.2, Student’s paired *t-*test). **k**, Pearson’s correlation coefficient *r* after infusion of saline or nAChR antagonist, during reward (*left*; blue), locomotion (*middle*; green), immobility (*right*; orange). Group means (± sem) shown in black. All comparisons vs. saline; Student’s paired *t*-test. **l**, Same as **k**, for DA-ACh coherence in 0.5-4 Hz band. **m**, Same as **k**, for DA-ACh phase offset. **n**, Experimental preparation for local pharmacological inhibition of iGluR and mAChR signaling in DLS. **o**, Average ACh fluorescence during immobility after infusion of saline, iGluR antagonists (p = 0.75 vs. saline, Student’s *t*-test) or mAChR antagonist (p = 0.004 vs. saline; Student’s *t*-test) as a percentage of pre-infusion signal. Note that blocking iGluRs does not significantly alter overall ACh levels compared to saline, suggesting that CINs continue to release ACh via spontaneous, cell-autonomous firing.

**Extended Data Fig. 6.**
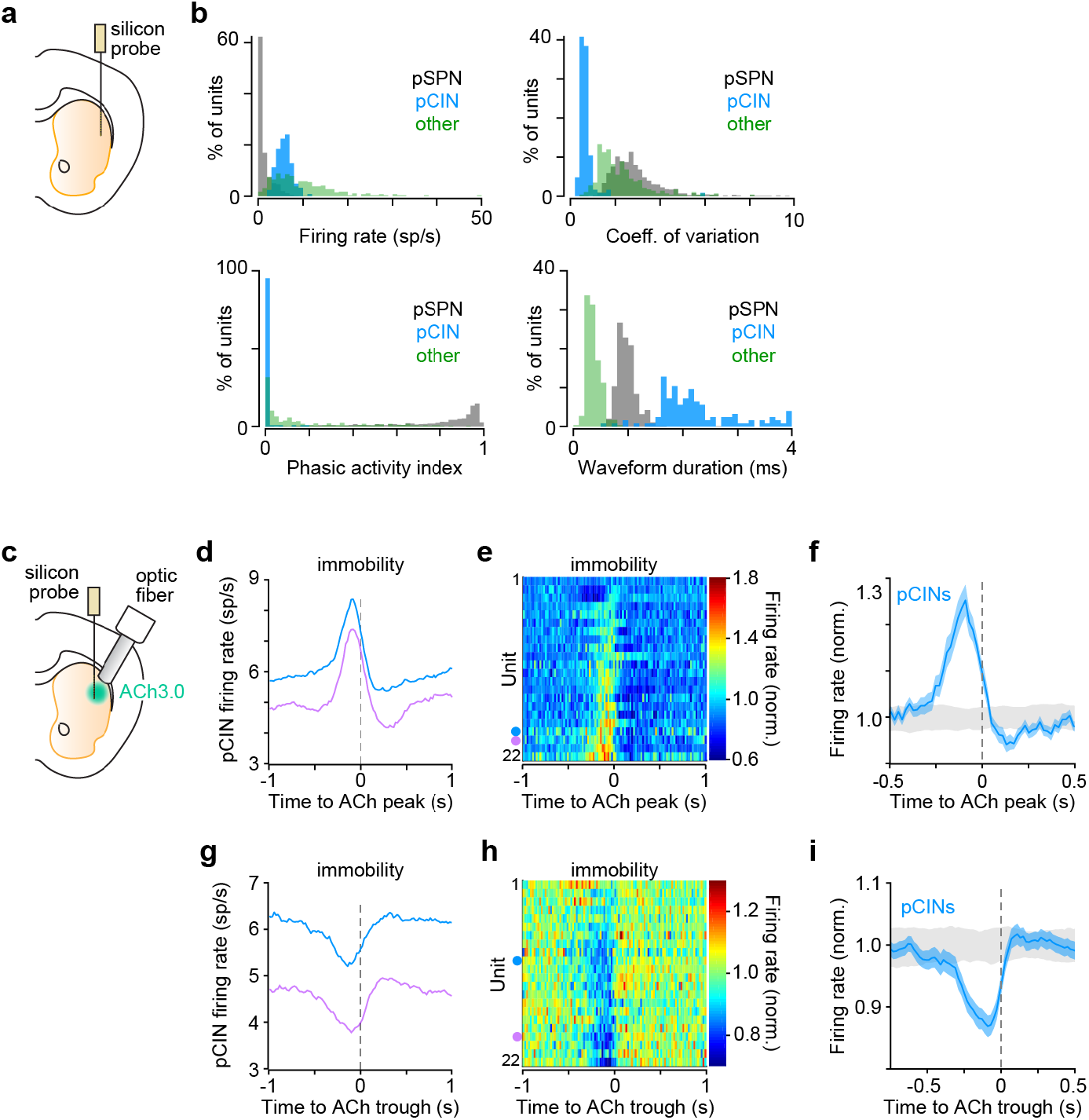
Periodic fluctuations in ACh are preceded by phasic changes in pCIN spiking. **a**, Experimental preparation for acute *in vivo* extracellular recordings in DLS. **b**, Distribution of firing rates (*top left*), coefficient of variation (*top right*), phasic activity index (*bottom left*), and waveform duration (*bottom right*) for pSPNs (black), pCINs (blue), and other interneurons (green). **c**, Experimental preparation for simultaneous ACh photometry and acute *in vivo* extracellular recordings in DLS. **d**, Firing rate of two example units aligned to ACh fluorescence peaks during immobility. **e**, Instantaneous firing rate of pCINs (n = 22; normalized to mean firing rate) aligned to ACh fluorescence peaks during immobility. Dots indicate units in **d. f**, Mean (± sem) firing rate of pCINs from **e**. 95% confidence interval shown in gray. **g**, Firing rate of same units as in **d** aligned to troughs in ACh fluorescence. **h**, Same as **e**, for firing rate of pCINs aligned to ACh fluorescence troughs. Dots indicate units in **g. i**, Mean (± sem) firing rate of pCINs from **h**. 95% confidence interval shown in gray.

**Extended Data Fig. 7.**
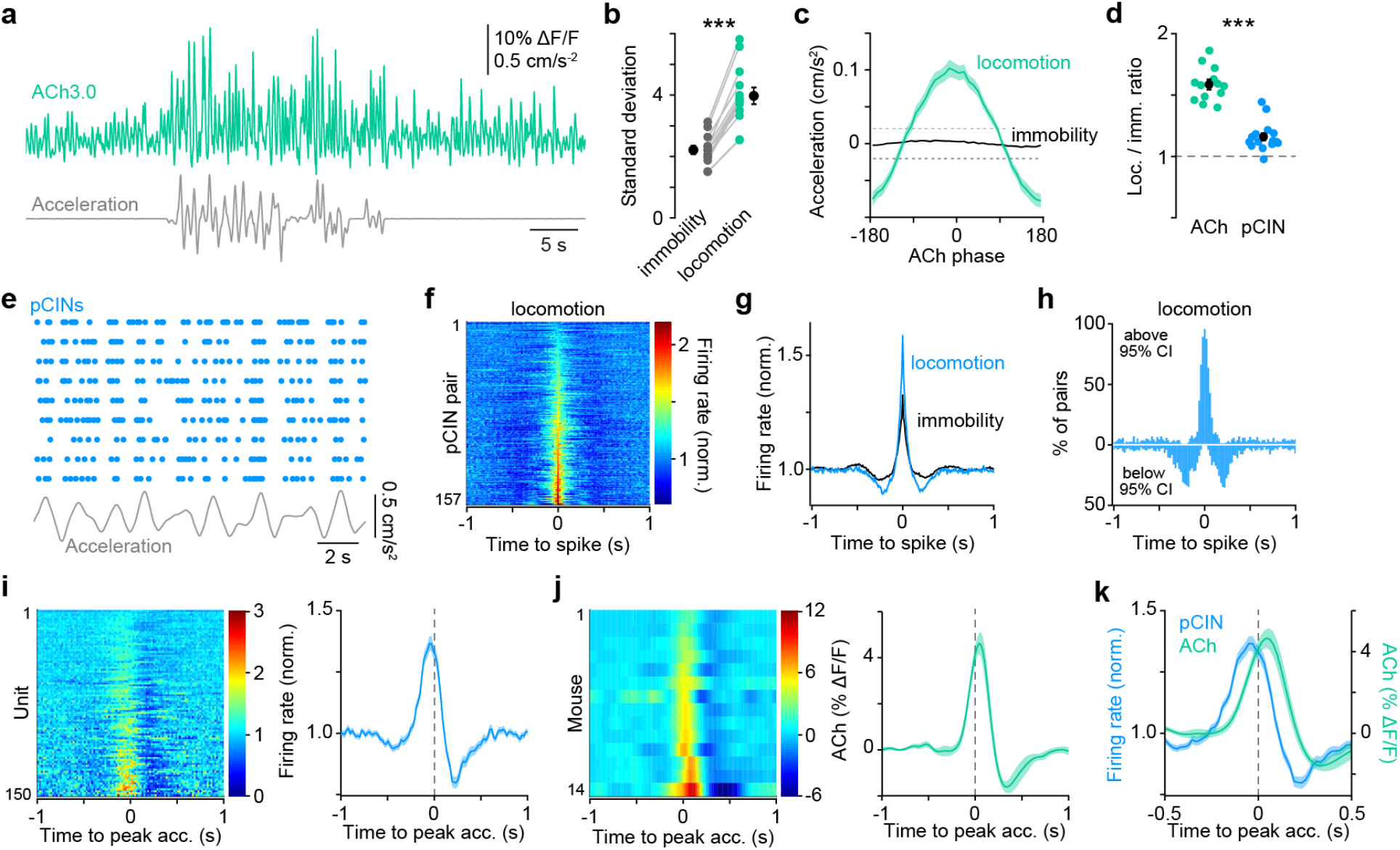
ACh transient amplitude and CIN coherence increase during locomotion. **a**, Continuous recording of ACh fluorescence (teal) and treadmill acceleration (gray). **b**, Standard deviation of ACh signal during immobility (black) and locomotion (teal). Group mean (± sem) shown in black (n = 14 mice; p = 2.06×10^−5^; Student’s *t*-test). **c**, Mean (± sem) acceleration at different phase of periodic ACh fluctuations in the 0.5-4 Hz frequency band during locomotion (teal) and immobility (black). Dashed lines depict 95% confidence interval. **d**, Fold change in mean ACh fluorescence (teal; n = 14 mice) and pCIN firing rate (blue; n = 15 mice) between periods of locomotion vs. immobility. Periodic fluctuations in ACh levels increase more notably than pCIN firing rates. **e**, Example spike raster for 9 simultaneously recorded pCINs (blue) and treadmill acceleration (gray) during a running bout on the treadmill. **f**, Heatmap of cross-correlograms (CCGs) for all pCIN-pCIN pairs (n = 157) normalized to baseline firing rate during locomotion. **g**, Mean (± sem) baseline-normalized CCG from all pCIN-pCIN pairs during locomotion (from **f**, in blue) and immobility (black) showing a near doubling in spike coherence. **h**, Proportion of CCGs from **f** exceeding 95% confidence interval (bin: 20 ms). **i**, *Left*, instantaneous firing rate of pCINs (n = 150 units) normalized to their mean rate aligned to acceleration peaks. *Right*, mean (± sem) firing rate from all units show at left. **j**, Same as **i** for ACh fluorescence (n = 14 mice). **k**, Overlay of peak acceleration-aligned pCIN discharge (blue) from **i** and ACh fluorescence (teal) from **j**. Mean (± sem) firing rate of pCINs (blue) and mean (± sem) ACh fluorescence aligned to positive acceleration peaks. pCIN spiking precedes ACh fluorescence by ∼100 ms.

**Extended Data Fig. 8.**
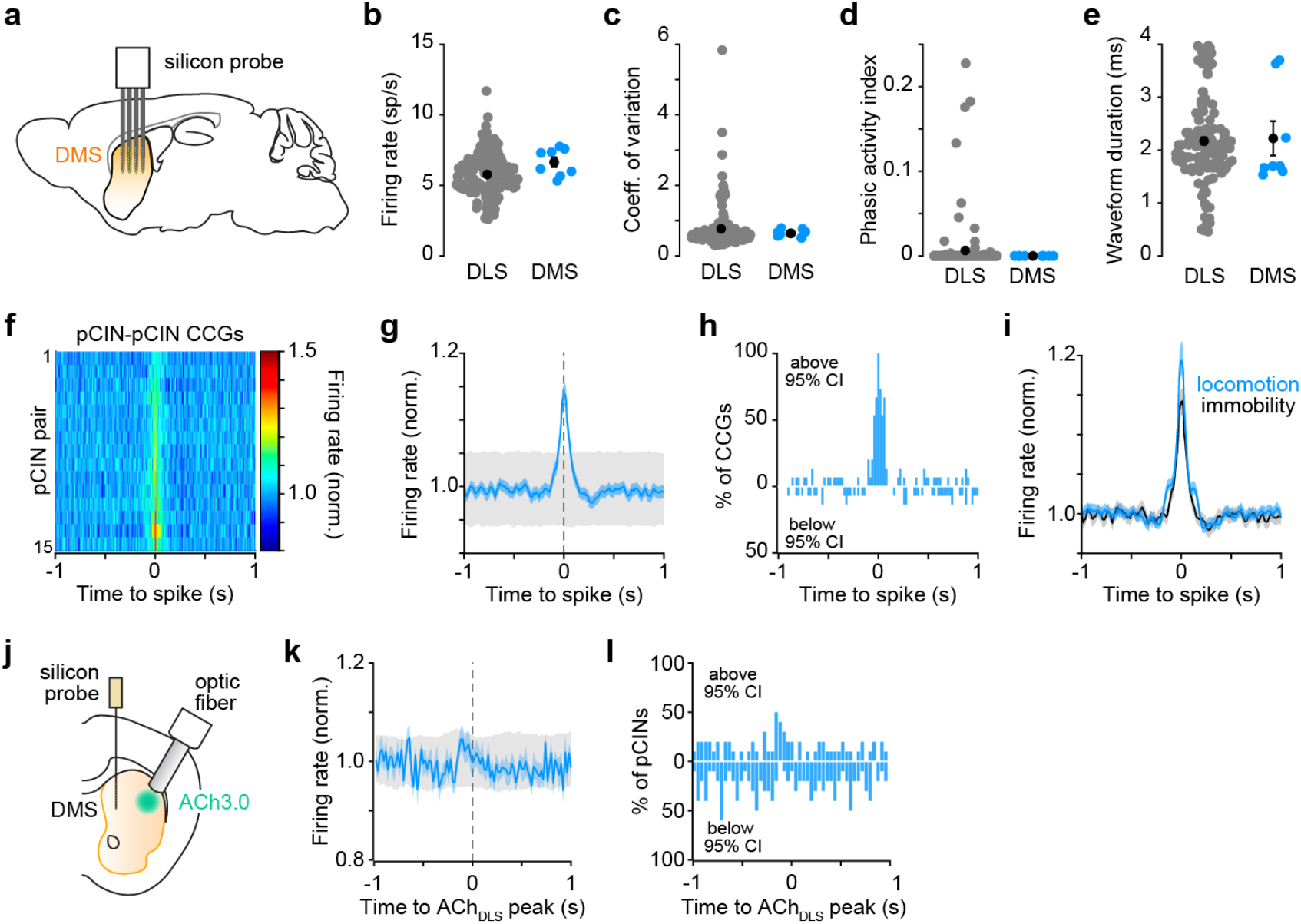
pCINs fire coherently with other pCINs in DMS, but not with ACh fluctuations in DLS. **a**, Diagram of experimental preparation. **b**, Comparison of discharge rate for pCINs recorded in DLS (black; n = 150) and DMS (blue; n = 8). Group means (± sem) shown in black (p = 0.06, Student’s two-sample *t*-test). **c**, Same as **b**, for coefficient of variation (p = 0.66, Student’s two-sample *t*-test). **d**, Same as **b**, for phasic activity index (p = 0.23, Student’s two-sample *t*-test). **e**, Same as **b**, for waveform duration (p = 0.48, Student’s two-sample *t*-test). **f**, Heatmap of cross-correlogram (CCGs) for all DMS pCIN-pCIN pairs normalized to mean firing rate during periods of immobility. **g**, Population mean (± sem) CCG from (F). 95% confidence interval shown in gray. **h**, Proportion of CCGs from **f** exceeding 95% confidence interval (bin: 20 ms). **i**, Same as **g** in black, overlaid with mean (± sem) CCG of pCIN-pCIN pairs in DMS during locomotor bouts (blue). **j**, Experimental preparation for acute *in vivo* extracellular recordings in DMS and ACh photometry in ipsilateral DLS. **k**, Mean (± sem) firing rate of DMS pCINs (n = 8) aligned to DLS ACh fluorescence peaks. 95% confidence interval shown in gray. **l**, Proportion of units from **k** exceeding 95% confidence interval (bin: 20 ms).

